# Developmental plasticity in visual cortex is necessary for normal visuomotor integration and visuomotor skill learning

**DOI:** 10.1101/2021.06.20.449148

**Authors:** Felix C. Widmer, Georg B. Keller

## Abstract

The experience of coupling between motor output and visual feedback is necessary for the development of visuomotor skills and shapes visuomotor integration in visual cortex. Whether these experience-dependent changes involve plasticity in visual cortex remains unclear. Here, we probed the role of NMDA receptor-dependent plasticity in mouse primary visual cortex (V1) during visuomotor development. Using a conditional knockout of NMDA receptors and a photoactivatable inhibitor of CaMKII, we locally perturbed plasticity in V1 during first visual experience, recorded neuronal activity in V1, and tested the mice in a visuomotor task. We found that perturbing plasticity before, but not after, first visuomotor experience reduces responses to unpredictable stimuli, diminishes the suppression of predictable feedback in V1, and impairs visuomotor skill learning later in life. Our results demonstrate that plasticity in the local V1 circuit during early life is critical for shaping visuomotor integration.

*** Dear reader, please note this manuscript is formatted in a standard submission format, and all statistical information is provided in Table S1. ***

## INTRODUCTION

Movement results in predictable sensory consequences. Through experience, the brain learns this mapping from motor output to sensory feedback. Raised without coupling between movements and visual feedback during visual development, kittens fail to use visual input to guide movements with the impairment specific to the dimension in which the artificial uncoupling occurred (Hein and Held, 1967; Held and Hein, 1963). The same coupling between locomotion and visual feedback is necessary to integrate visual and motor-related signals in primary visual cortex (V1). Under normal conditions, V1 has been shown to exhibit distinct and salient responses to unpredictable mismatches between movement and visual feedback in both humans and mice (Keller et al., 2012; Stanley and Miall, 2007; Zmarz and Keller, 2016). In mice raised from birth without coupling between movement and visual feedback, mismatch responses are absent and only emerge after first exposure to normal visuomotor coupling (Attinger et al., 2017). Thus, the coupling between movement and visual feedback is essential for both visuomotor behavior and normal visuomotor integration in V1. It is still unclear, however, where the plasticity occurs that is driven by the experience of coupling between movement and visual feedback.

Given that V1 receives both the bottom-up visual input and signals consistent with a top-down prediction of visual feedback given movement (Leinweber et al., 2017) necessary to compute these mismatch responses, it has been speculated that mismatch responses are computed locally in V1. Neurons in layer 2/3 (L2/3) of V1 that are responsive to visuomotor mismatch receive balanced and opposing top-down motor-related and bottom-up visual input (Jordan and Keller, 2020), consistent with a subtractive computation of mismatch responses. Thus, it is possible that visuomotor experience establishes this balance between top-down and bottom-up input on individual L2/3 neurons in V1. If this were so, we would predict that the consequence of perturbing plasticity locally in V1 during visuomotor development would be reduced mismatch responses in these neurons.

Here, we tested this by interfering with plasticity locally in V1 during first visuomotor experience using two separate approaches. First, we used a local knockout of N-Methyl-D-aspartate (NMDA) receptors in V1 prior to first visuomotor experience. NMDA receptors are known to be involved in a wide variety of different forms of plasticity (Paoletti et al., 2013; Rodriguez et al., 2019), and are necessary for activity-dependent synaptic strengthening in cortex (Hasan et al., 2013; Kirkwood and Bear, 1994; Lo et al., 2013). In a parallel approach, to impair plasticity in V1 in a cell type specific manner, we used a photoactivatable inhibitor of the calcium/calmodulin-dependent protein kinase II (CaMKII). CaMKII has been shown to be an essential element of NMDA receptor-dependent plasticity (Barria and Malinow, 2005; Gambrill and Barria, 2011; Wang et al., 2011). NMDA receptors are thought to exert their influence on synaptic plasticity by increasing calcium influx into the cell, where calmodulin binds calcium and activates CaMKII. The idea that NMDA receptors and CaMKII are on the same plasticity pathway is supported by a number of findings. For example, spine enlargement triggered by NMDA receptor stimulation can be inhibited by blocking CaMKII (Herring and Nicoll, 2016). Additionally, activated CaMKII and NMDA receptors directly interact (Leonard et al., 1999) to form CaMKII-NMDA receptor complexes that are required for the induction of long-term potentiation (Barria and Malinow, 2005), and likely control synaptic strength (Lisman et al., 2012). Thus, we expected that NMDA receptor knockout and CaMKII inhibition would have similar effects on the responses of L2/3 neurons. We find that both types of manipulations systematically impair the development of normal visuomotor integration in L2/3 neurons, commensurate with the impairment observed in mice that are raised without experience of the coupling between movement and visual feedback (Attinger et al., 2017), but differ in the way they influence visual responses.

## RESULTS

### Knockout of *Grin1* prior to first visual experience impaired the development of normal visual and visuomotor mismatch responses

To determine the dependence of visuomotor integration specifically on local plasticity in V1 during development, we quantified the effect of a conditional knockout of NMDA receptors in V1, prior to first visual experience, on functional responses in V1 L2/3 neurons. To achieve this, we used NR1^flox^ mice, which carry a modified version of the *Grin1* gene (also referred to as *NMDAR1*, an essential subunit of the NMDA receptor) that can be rendered inactive by Cre recombination (Tsien et al., 1996). We dark reared these mice from birth and injected an adeno-associated viral vector (AAV2/1-EF1α-Cre-T2A-mCherry) to express Cre recombinase unilaterally into V1 at postnatal day P21, prior to first visual experience (*ΔGrin1*_juv_; **Figures 1A and 1B**). At P30 we then injected a second AAV vector to express GCaMP6f (AAV2/1-EF1α-GCaMP6f) bilaterally in both primary visual cortices to record neuronal activity in the knockout hemisphere and a within-mouse control hemisphere. Mice were then exposed to visual input for the first time in their life at P32, when they were introduced to a virtual environment that provided closed-loop feedback between forward locomotion and backward visual flow in a virtual corridor (Attinger et al., 2017). Mice were trained in this setup for 2 hours every other day for 12 days (for a total of 6 sessions), after which we then measured calcium activity in L2/3 neurons using two-photon imaging (**Figure 1C**). We validated the method for the local knockout of *Grin1*, using an in situ hybridization with a *Grin1* mRNA probe in a subset of mice and found a marked reduction at the injection site of the Cre vector (**Figure 1D**). During the imaging experiments, mice were first exposed to closed-loop visual flow feedback in a virtual corridor (see Methods). To measure mismatch responses, we introduced brief (1 s) halts of visual flow at random times (Keller et al., 2012). To estimate the contributions of visual flow and locomotion separately, mice were then presented with a playback of the visual flow they had previously self-generated in the closed-loop session (we will refer to this as the open-loop session). To measure visual responses, mice were presented with full-field drifting gratings of different orientations. Finally, to isolate motor-related signals, we measured locomotion-related activity in complete darkness.

**Figure 1.**
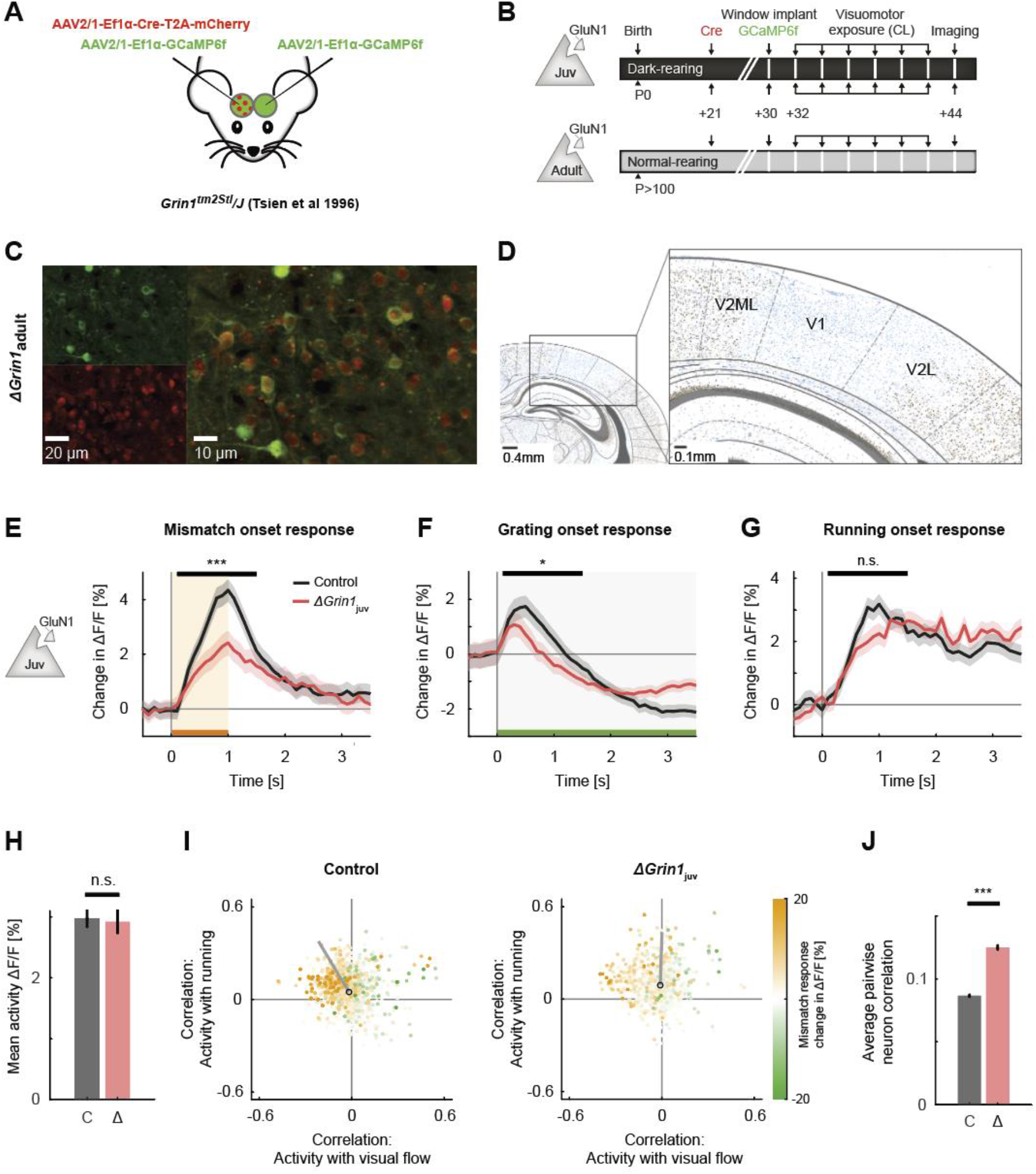
NMDA receptor knockout prior to first visual experience impaired the development of normal visual and mismatch responses. **(A)** We injected an AAV to express Cre recombinase unilaterally and another to express a calcium indicator bilaterally (GCaMP6f) in V1 of *ΔGrin1* mice prior to first visual experience. **(B)** Experimental timeline: a first group of mice (*ΔGrin1_juv_*) was dark reared from birth. We injected an AAV to express Cre at P21 unilaterally in V1, injected a second AAV to express GCaMP6f bilaterally, and implanted imaging windows bilaterally at P30. A second group of mice (*ΔGrin1_adult_*) was reared normally and received the same injections at P>100. All mice then had 6 sessions of visuomotor exposure in a closed-loop (CL) virtual environment before imaging experiments. **(C)** Example two-photon images showing co-expression of GCaMP6f and Cre-mCherry constructs. **(D)** In situ hybridization against *Grin1* mRNA (see Methods) confirming the local knockout of *Grin1* in V1. Blue: Hematoxylin stain for cell nuclei; brown: *Grin1* hybridization signal. Brain regions were identified using a mouse brain atlas (Franklin and Paxinos, 2012). **(E)** The average population response (stimulus induced change in ΔF/F) to mismatch was stronger in control (black) than in *ΔGrin1*_juv_ (red) hemispheres. Shading indicates standard error of the mean (SEM) across neurons. Orange shading and bar indicate duration of mismatch. Mean responses were compared across neurons in the time window indicated by the black bar above the traces. Here and in subsequent panels, n.s.: p > 0.05, *: p < 0.05, **: p < 0.01, ***: p < 0.001. For all details of statistical testing, see **Table S1**. **(F)** As in **E**, but for drifting grating responses (see Methods). Green shading and bar indicate presence of a grating stimulus. **(G)** As in **E**, but for running onset responses in closed-loop sessions. **(H)** Mean calcium activity of neurons in the control (C, gray) and *ΔGrin1*_juv_ (Δ, red) hemisphere during the closed-loop session. Error bars indicate SEM across neurons. **(I)** Scatter plot of the correlation between neuronal activity and visual flow, and the correlation between neuronal activity and running speed in open-loop sessions for all L2/3 neurons recorded in control (left) and *ΔGrin1*_juv_ (right) hemispheres. Each dot shows the correlations for one neuron, and dot color indicates the neuron’s mismatch response. Black circles mark the population mean, and solid black lines indicate the direction of the first principal component of the distribution (see **Figure S1B** and Methods). **(J)** Average pairwise correlation of neuronal activity was higher in *ΔGrin1*_juv_ (Δ, red) compared to that in the control (C, black) hemisphere. Error bars indicate SEM across neurons.

We found that visuomotor mismatch responses in the knockout hemisphere were reduced compared to the control hemisphere (**Figure 1E**), commensurate with the response reduction in mice that never experienced coupling between locomotion and backward visual flow (**Figure S1A**). We also found a reduction in grating onset responses (**Figure 1F**), but no evidence of a reduction in motor-related activity upon running onset in a closed-loop environment (**Figure 1G**). The fact that mismatch and visual responses are influenced by NMDA receptor knockout is consistent with impairment of the comparator function of L2/3 (Jordan and Keller, 2020). An alternative explanation would be that the reduced responses are simply a consequence of an overall reduction in activity levels. However, this was not the case as comparing mean activity levels between control and knockout hemisphere showed no evidence of a reduction in activity (**Figure 1H**). Mismatch responses are thought to arise from a transient imbalance between opposing bottom-up visual inhibition and top-down motor-related excitation. A reduction of mismatch responses could be the result of a reduction in either top-down or bottom-up input, or a failure to appropriately match bottom-up inhibition and top-down excitation. To disambiguate these two possibilities, we estimated the contribution of bottom-up visual input and top-down motor-related input by calculating the correlation between neuronal activity and visual flow, and that between neuronal activity and locomotion for each neuron (**Figure 1I**). Consistent with responses in mice without an NMDA receptor knockout (Attinger et al., 2017), we found that in the control hemisphere neurons with high mismatch responses tended to show a negative correlation with visual flow and a positive correlation with running speed. In the knockout hemisphere we found that both the average correlation of activity with running speed and correlation of activity with visual flow were slightly, but significantly increased relative to the control hemisphere (mean correlation of activity with visual flow control hemisphere: −0.017, *ΔGrin1*_juv_ hemisphere: −0.010, p < 10^−5^; mean correlation of activity with running speed control hemisphere: 0.048, *ΔGrin1*_juv_ hemisphere: 0.090, p < 10^−5^; two-sample independent t-test). The overall distribution resembled the distribution we had observed previously in mice raised without coupling between running and visual flow (Attinger et al., 2017). We quantified this using the angle of the first principal component of the distribution relative to the axis defined by the correlation of activity with running speed. Similar to mice raised with coupling between running and visual flow, we found that in the control hemisphere the majority of neurons exhibited opposing signs of correlation with running and visual flow, which manifested as a principal component close to the negative diagonal. In the knockout hemisphere the distribution is shifted in the direction of that observed in mice raised without coupling between running and visual flow, where the principal component is rotated towards the positive diagonal (**Figure S1B**). These results are consistent with the interpretation that the NMDA receptor knockout interferes with the establishment of the balance between opposing top-down and bottom-up input in individual neurons. Lastly, consistent with the effect of systemic inhibition of NMDA receptors on correlations of L2/3 neurons in V1 (**Figure S1D**) (Hamm et al., 2017), we found that in the knockout hemisphere the average pairwise correlation of neuronal activity was higher compared to that in the control hemisphere (**Figure 1J**). Thus, NMDA receptor knockout prior to first visual experience prevents the development of normal visual and visuomotor mismatch responses in V1.

These results would be consistent with either a role of the NMDA receptor in the plasticity necessary for the establishment of visuomotor integration in V1, or a direct involvement of NMDA receptors in generating neuronal calcium responses. The latter could be driven by an influence of NMDA receptors on the overall excitability of the neurons, or, given that NMDA receptors conduct calcium, by directly reducing the calcium response. To disambiguate this, we repeated the same NMDA receptor knockout experiments in a second group of mice that had been reared in a normal light-dark cycle (*ΔGrin1*_adult_; **Figure 1B**). We found that in these mice there was no difference in any of the responses between those in the control hemisphere and those in the knockout hemisphere (**Figures 2A-2C**). Consistent with the finding that pharmacological inhibition of NMDA receptors in adult mice results in an overall decrease of V1 activity (Ranson et al., 2019) (**Figure S1C**), we found a strong reduction in overall activity levels in the knockout hemisphere (**Figure 2D**). Consistent with a lack of an NMDA receptor knockout induced change in mismatch and visual responses, the distribution of visual flow and running correlations with activity in control and knockout hemispheres was very similar (**Figure 2E**). Lastly, as in the juvenile knockout, we found an increase in the average correlation between neurons (**Figure 2F**). This increase in correlation is likely specific to L2/3 neurons, as a similar knockout in layer 4 (L4) neurons results in a decrease in correlation between neurons that lack NMDA receptors (Mizuno et al., 2021). This demonstrates that NMDA receptors are not necessary to maintain mismatch and visual responses once V1 is fully trained by visuomotor experience.

**Figure 2.**
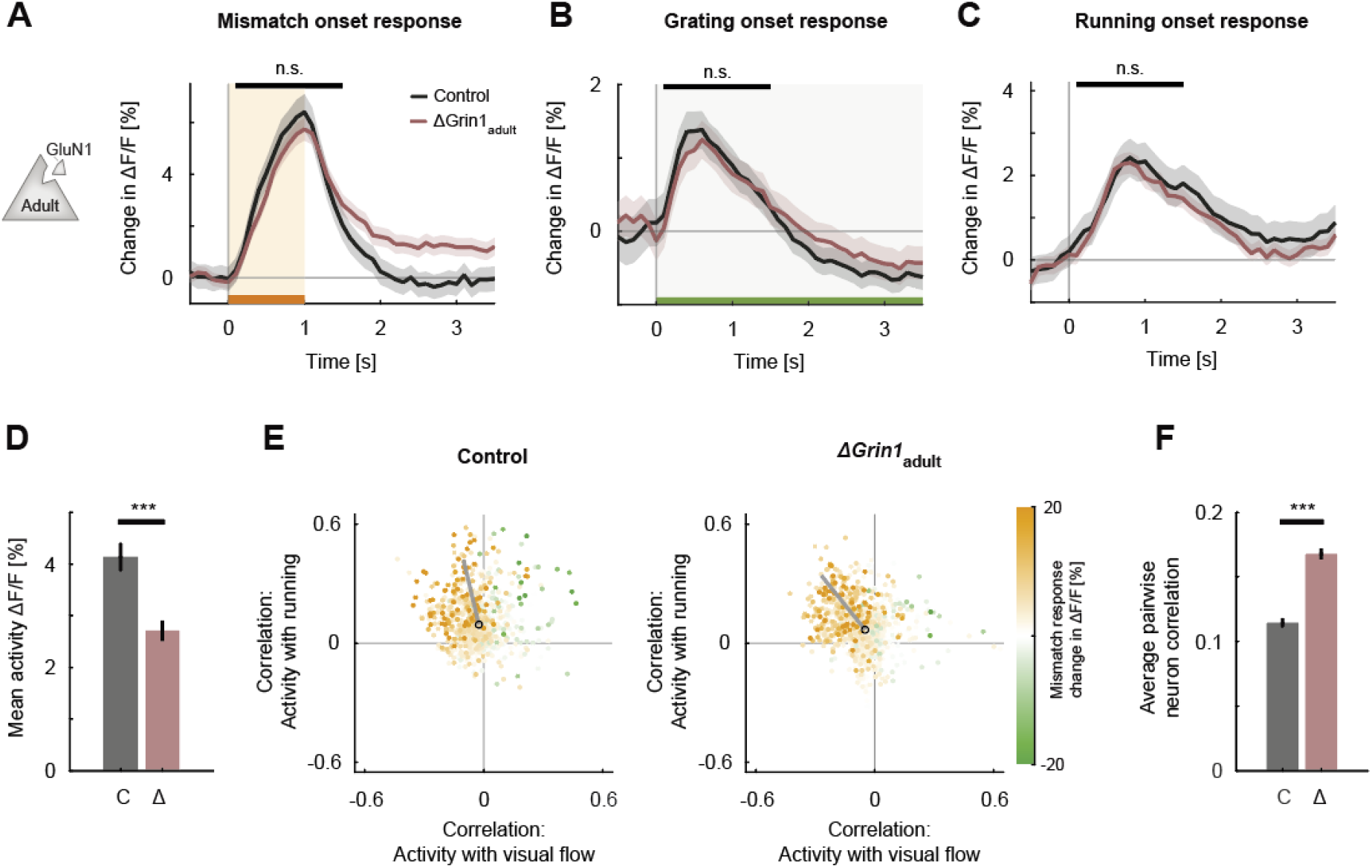
NMDA receptor knockout in the adult mouse did not impair visual and visuomotor responses. **(A)** The average population response to mismatch was similar in control (black) and in *ΔGrin1*_adult_ (dark red) hemispheres. Shading indicates SEM across neurons. Orange shading and bar indicate duration of mismatch. Mean responses were compared across neurons in the time window indicated by the black bar above the traces. Here and in subsequent panels, n.s.: p > 0.05, *: p < 0.05, **: p < 0.01, ***: p < 0.001. For all details of statistical testing, see **Table S1**. **(B)** As in **A**, but for responses to the onset of a drifting grating stimulus (see Methods). Green shading and bar indicate presence of grating stimulus. **(C)** As in **A**, but for running onset responses in closed-loop sessions. **(D)** Mean activity of neurons in the control (C, gray) and *ΔGrin1*_adult_ (Δ, dark red) hemisphere during the closed-loop session. Error bars indicate SEM across neurons. **(E)** Scatter plot of the correlation between neuronal activity and visual flow, and the correlation between neuronal activity and running speed in open-loop sessions for all L2/3 neurons recorded in in control (left) and *ΔGrin1*_adult_ (right) hemispheres. Each dot shows the correlations for one neuron, and dot color indicates the neuron’s mismatch response. Black circles mark the population mean, and solid black lines indicate the direction of the first principal component of the distribution (see Methods). **(F)** Average pairwise correlation of neuronal activity was higher in *ΔGrin1*_adult_ (Δ, dark red) compared to that in the control (C, black) hemisphere. Error bars indicate SEM across neurons.

Both a visuomotor mismatch and the sudden appearance of a visual stimulus are unpredictable events and can be interpreted as negative and positive prediction errors, respectively. Assuming there is indeed a deficit in the development of prediction error responses induced by the NMDA receptor knockout, we would also expect a similar deficit in the suppression of predictable responses. To investigate this, we quantified the suppression of running onset responses by visual flow in the closed-loop session. In normally reared mice, a running onset with closed-loop visual feedback is typically associated with an increase in activity that is transient, whereas the response to the same running onset in darkness results in a sustained change in mean activity (**Figure 3A**). One interpretation of this is that the visual flow coupled to locomotion in the closed-loop session triggers a suppression of the running-related responses. We can quantify the suppression in the closed-loop session by taking the difference between the running onset response in darkness and that in the closed-loop session (**Figure 3A**). Computing this difference for control mice, *ΔGrin1*_juv_ mice, and *ΔGrin1*_adult_ mice, we found that this suppression was absent only in the knockout hemisphere of the *ΔGrin1*_juv_ mice (**Figures 3B and 3C**). This is consistent with an impairment in the suppression of predictable responses in L2/3 neurons by an NMDA receptor knockout prior to first visual experience.

**Figure 3.**
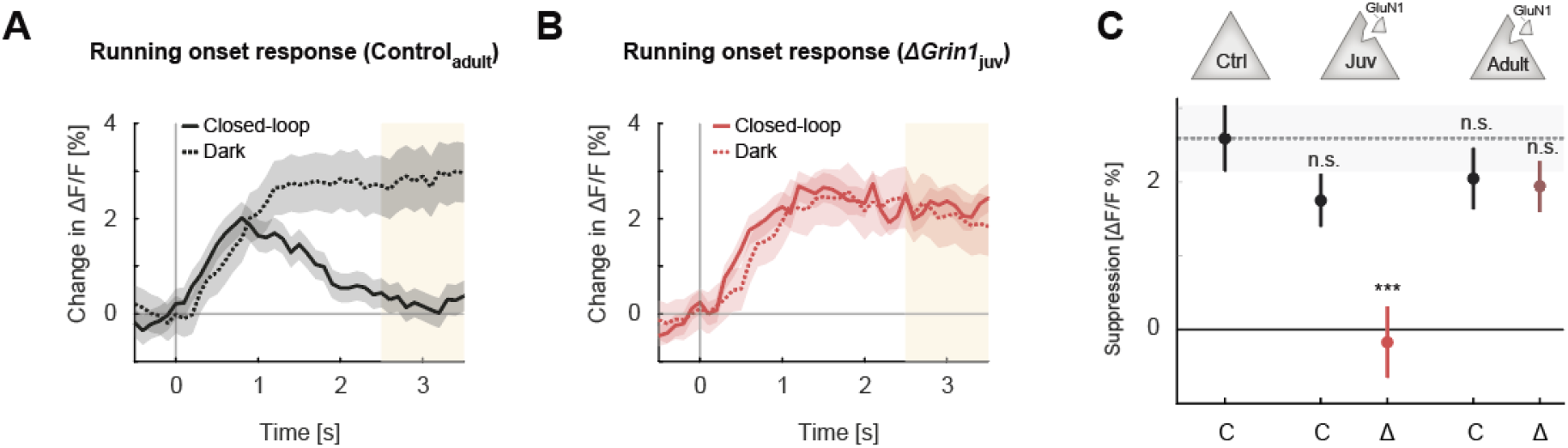
Suppression of running onset responses by visual flow was reduced by an NMDA receptor knockout prior to first visual experience. **(A)** The average population response to running onset in closed-loop sessions (solid) and dark sessions (dotted) in adult control mice. Shading indicates SEM across neurons. Albescent white shading marks analysis window used in **C**. Note, the visual flow associated with closed-loop running results in a suppression of motor-related responses. **(B)** As in **A**, but for *ΔGrin1*_juv_ data in the knockout hemisphere. **(C)** Average closed-loop visual feedback induced suppression of activity for all neurons in adult control mice and control (C) or knockout (Δ) hemispheres of *ΔGrin1*_juv_ and *ΔGrin1*_adult_ mice. Suppression is calculated as the difference between the running onset response in dark and closed-loop session in the window 2.5 s to 3.5 s after running onset, marked in **A**and **B**. Error bars indicate SEM across neurons. Comparison against data from control mice; n.s.: p > 0.05, ***: p < 0.001. For all details of statistical testing, see **Table S1**.

### Local NMDA receptor dysfunction during development resulted in impaired visuomotor skill learning later in life

Assuming developmental plasticity is necessary for the establishment of normal visuomotor integration, we expected that the *ΔGrin1*_juv_ mice would exhibit behavioral impairments in cortex-dependent visuomotor tasks. To test this, we trained 6 *ΔGrin1*_juv_ mice in a visuomotor task later in life. For these experiments we used two control groups. The first was composed of 13 *ΔGrin1*_adult_ mice, and the second was composed of 6 control mice (Control_juv_) that did not receive a *Grin1* knockout but were dark reared from birth. The *ΔGrin1*_juv_ and Control_juv_ groups were dark reared until P32. All three groups were initially exposed to closed-loop experience in a virtual reality setup as described above and subsequently trained to perform a virtual navigation task (Heindorf et al., 2018) (**Figures 4A and 4B**). In this task, mice had control over movement in a virtual environment through rotation and forward locomotion on a spherical treadmill and were trained to reach the end of a virtual corridor for a water reward. Training lasted for 7 days, 1 hour per day. We quantified performance using an index that is based on the fraction of distance traveled toward the target normalized by the total distance traveled (see Methods). The dark reared Control_juv_ mice and the adult knockout *ΔGrin1*_adult_ mice both learned to perform the task over the course of the training. The *ΔGrin1*_juv_ mice, however, failed to show evidence of increased performance over the course of the 7 days of training, and exhibited significantly reduced performance compared to the two control groups late in training (**Figure 4C**). To test for the mice’s ability to induce a behavioral response to an unexpected perturbation of visual feedback, we introduced sudden offsets of the current heading direction at random times by 30° either to the left or to the right. With training, mice learned to correct for these offset perturbations with a turn that corrected for the offset. Both Control_juv_ and *ΔGrin1*_adult_ mice corrected for offset perturbations with a compensatory turn in the correct direction by the end of training (**Figure 4D**). However, the *ΔGrin1*_juv_ mice failed to correct for these offsets. Quantifying this as the learning related change in offset perturbation response, we found that Control_juv_ and *ΔGrin1*_adult_ mice both exhibit larger learning related changes than the *ΔGrin1*_juv_ mice (**Figure 4E**). Thus, consistent with the dependence of normal visuomotor integration on NMDA receptors during first visuomotor experience, we found that mice that lack NMDA receptors during first visuomotor experience are impaired in learning certain cortex-dependent visually guided motor tasks later in life.

**Figure 4.**
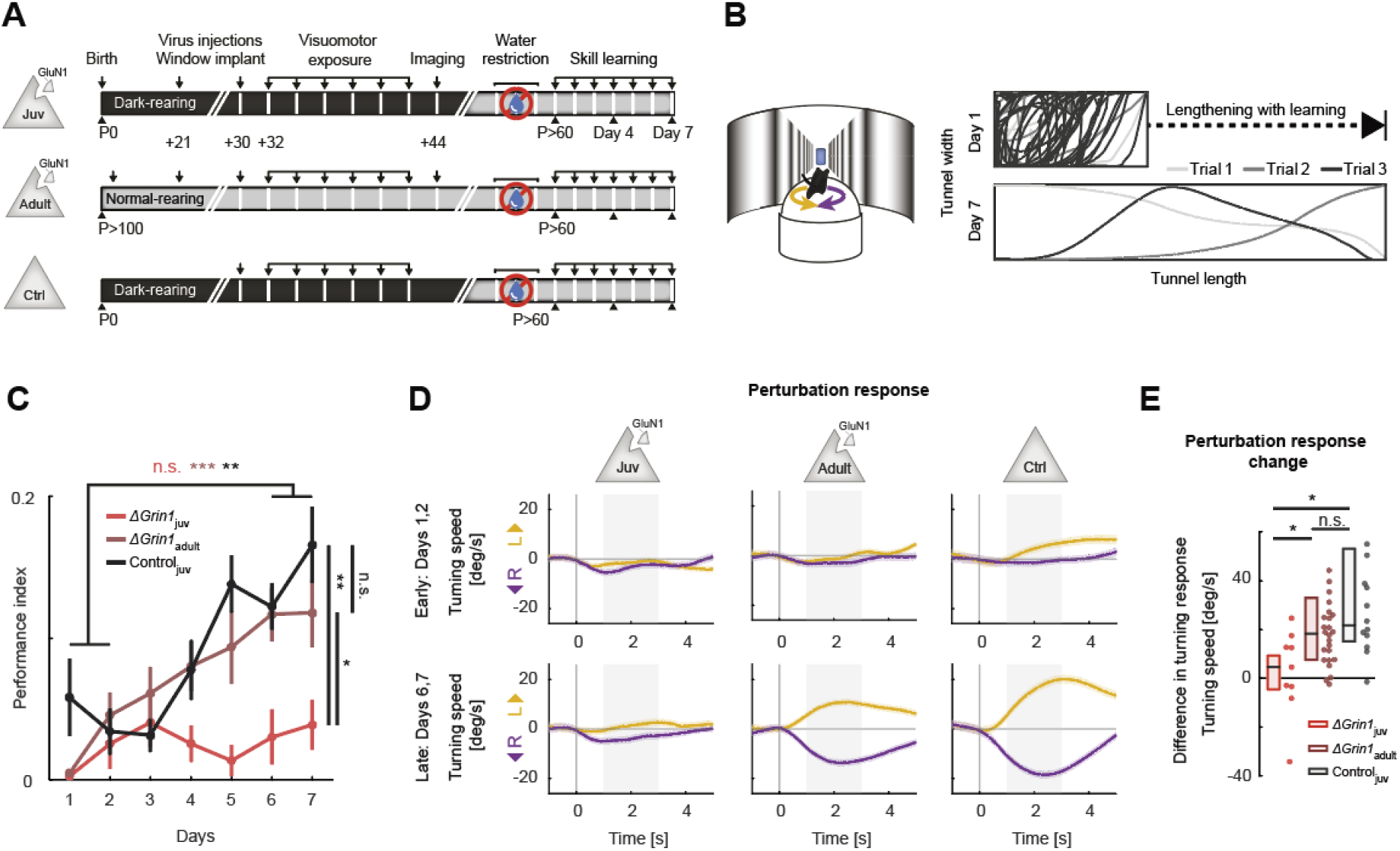
NMDA receptor knockout in V1 before first visuomotor experience impaired learning of a visuomotor task later in life. (A) Experimental approach and timeline. Three groups of mice were trained: the first was composed of 6 *ΔGrin1*_juv_ dark reared mice, the second was composed of 13 *ΔGrin1*_adult_ normally reared mice, and the third was composed of 6 C57/Bl6 dark reared control mice. Mice were water restricted and subsequently trained to perform a virtual navigation task (see Methods). **(B)** Left: Schematic of virtual reality setup. Mice controlled forward translational motion and rotation in a virtual corridor by rotating a spherical treadmill and were trained to navigate to the end of a corridor for a water reward. As performance increased, the task difficulty was increased by lengthening the virtual corridor. Right: Top-down view of the virtual corridor showing the trajectories of the mouse in three example trials (different gray levels) on day 1 (top) and day 7 (bottom). The ratio of virtual corridor length to width is not to scale. **(C)** Task performance as a function of training day (see Methods) of *ΔGrin1*_juv_ mice (red), *ΔGrin1*_adult_ mice (dark red), and dark reared control mice (Control_juv_, black) over the course of 7 days. Error bars indicate SEM across mice. *ΔGrin1*_adult_ and Control_juv_ mice exhibited performance improvements over the course of training, while *ΔGrin1*_juv_ mice did not (comparing average performance on day 1 and 2 (early), vs average performance on day 6 and 7 (late)). Performance on day 7 was different between *ΔGrin1*_juv_ and both *ΔGrin1*_adult_ and Control_juv_ mice. Here and in subsequent panels, n.s.: p > 0.05, *: p < 0.05, **: p < 0.01, ***: p < 0.001. For all details of statistical testing, see **Table S1**. **(D)** Turning response behavior (rotational velocity) following a sudden heading displacement of 30° (perturbation) to the left (yellow) or right (purple) of *ΔGrin1*_juv_, *ΔGrin1*_adult_ and Control_juv_ mice, early (top row) and late (bottom row) in training. Shading indicates SEM across trials. Gray shading indicates analysis window (+1 s to +3 s) used for quantification in **E**. **(E)** Quantification of perturbation offset responses shown in **D** as the difference between average left and right perturbation turning responses, late (bottom row in **D**) minus early (top row in **D**) in training. Boxes show median and quartiles, all data (averages over mice) are shown as dots to the right. *ΔGrin1*_adult_ and Control_juv_ mice learned to initiate corrective turns in response to visual offset perturbations, while *ΔGrin1*_juv_ mice did not.

### CaMKII-dependent plasticity in SST interneurons was necessary for feed-forward visual inhibition

Central to the subtractive computation of prediction error responses are inhibitory interneurons. By implementing the opposing influence of visual and locomotion related input in L2/3 neurons (Jordan and Keller, 2020), they allow for a subtraction of a bottom-up sensory input and a top-down prediction to compute prediction errors (Keller and Mrsic-Flogel, 2018). Based on measurements of calcium responses to visuomotor mismatches and artificial manipulations of activity in different interneuron subtypes, we have previously speculated that a subset of somatostatin (SST) positive interneurons mediates the visually driven inhibition necessary for negative prediction error responses in V1 L2/3 excitatory neurons (Attinger et al., 2017). We thus set out to test whether an impairment of plasticity selectively in SST interneurons in V1 during first visuomotor experience would result in a failure to establish visually driven inhibition in L2/3 neurons. To do this, we turned to a method that would allow us to target the intervention to SST interneurons selectively in V1. We used a photoactivatable autocamtide inhibitory peptide 2 (paAIP2) (Murakoshi et al., 2017) to inhibit calcium/calmodulin-dependent kinase II (CaMKII) using blue light illumination.

We repeated the experiments we performed with the NMDA receptor knockout (**Figures 1–3**) using paAIP2 in three groups of mice to target CaMKII inhibition either to excitatory neurons, SST interneurons (**Figure 5A**), or parvalbumin (PV) positive interneurons (**Figure S3A**). Again, all mice were dark reared from birth and received 2 hours of visuomotor experience in virtual reality environment every other day for 12 days (**Figures 5B and S3B**). The first group consisted of 6 C57/Bl6 mice that received an injection of an AAV to express paAIP2 under a CaMKIIα(1.3kb)-promoter (AAV2/1-CaMKIIα-mEGFP-P2A-paAIP2) in right V1. The other two groups consisted of 7 SST-Cre mice and 6 PV-Cre mice that each received an injection of AAV2/1-DIO-mEGFP-P2A-paAIP2 unilaterally in V1. At P30, prior to first visuomotor experience, mice were injected with an AAV to express a red-shifted calcium indicator (AAV2/1-Ef1α-NES-jRGECO1a) in both visual cortices. To activate paAIP2 during visuomotor exposure while mice were on the virtual reality setup, we illuminated V1 bilaterally using a blue (473 nm) laser, through the glass windows implanted for subsequent two-photon imaging (see Methods). As before (**Figures 1E-1G** and **Figures 2A-2C**), we then proceeded to measure mismatch, grating, and running onset responses in L2/3 neurons at P44. Similar to the responses observed in *ΔGrin1*_juv_ mice, we found that with paAIP2 expressed under the CaMKIIα(1.3kb)-promoter, the strongest changes were in mismatch and visual responses, while running onset responses were less affected (**Figures 5C-5E**). Mismatch responses were again reduced in the inhibited hemisphere compared to the control hemisphere (**Figure 5C**). Intriguingly, CaMKII inhibition resulted in a massive increase in visually driven activity of L2/3 neurons (**Figure 5D**). We speculate that this difference can be explained by the fact that the power of light used to activate paAIP2 falls off exponentially with cortical depth (**Figure S2A**) and that CaMKII inhibition predominantly influences superficial synapses, which preferentially carry top-down signals (see Discussion). Despite the difference in the effect of the manipulation on visually driven responses, the effect of the CaMKII inhibition on correlations between the activity of L2/3 neurons was an increase, similar to that observed in the *ΔGrin1*_juv_ mice (**Figure S2B**).

**Figure 5.**
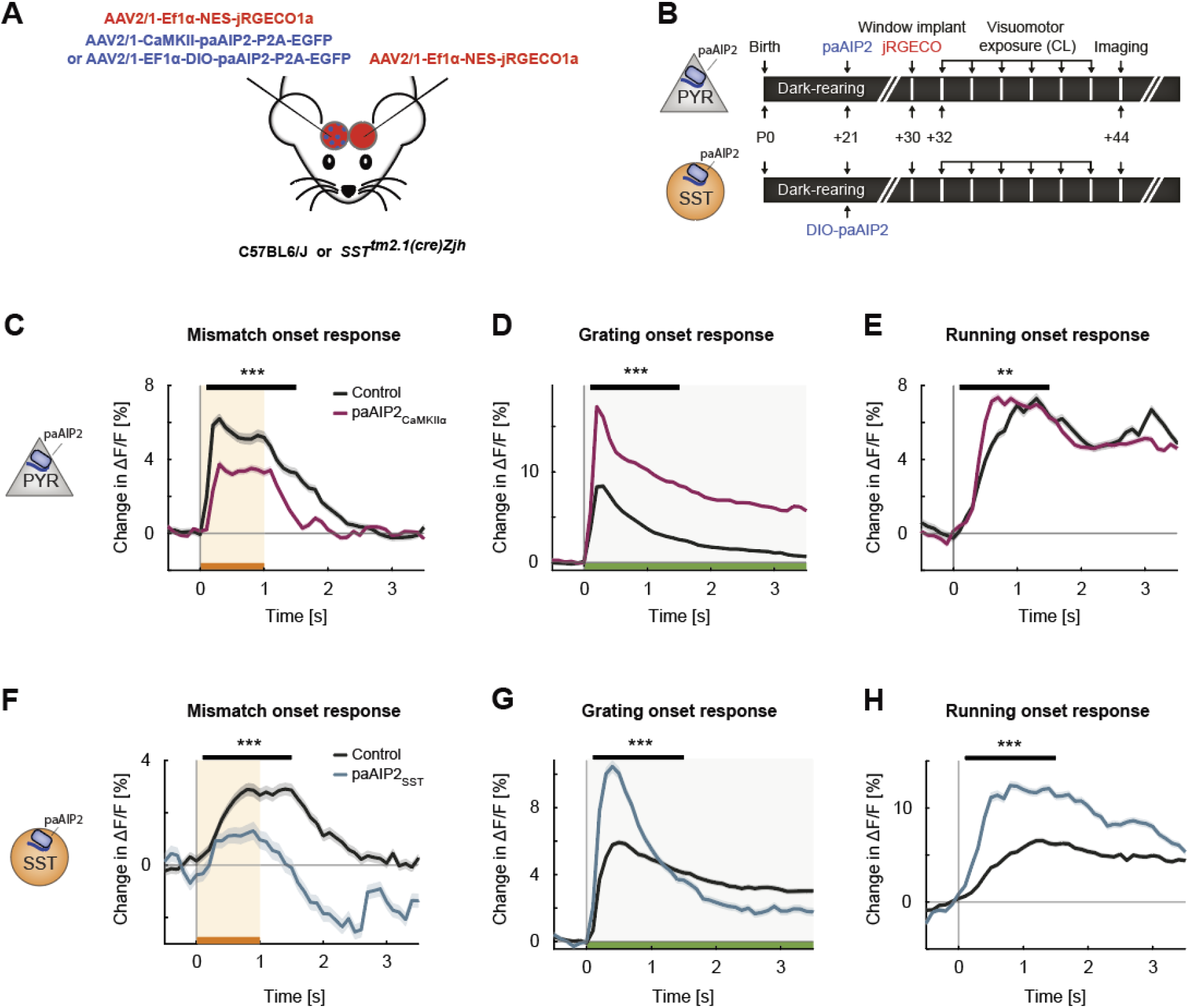
Inhibiting CaMKII in excitatory neurons or SST interneurons resulted in imbalanced visuomotor responses in L2/3 excitatory neurons. **(A)** We unilaterally injected an eGFP-tagged paAIP2 or DIO-paAIP2 expressing virus and a calcium indicator (jRGECO1a) bilaterally in V1. **(B)** Mice were dark reared from birth. AAV injections occurred at postnatal day 21 (to deliver paAIP2 or DIO-paAIP2) and P30 (to deliver jRGECO1a). Imaging window implantation occurred on P30. Mice had 6 sessions of visuomotor exposure in a closed-loop virtual environment during which we illuminated cortex bilaterally with blue light (473 nm) to inhibit CaMKII. We used of 6 C57/Bl6J mice, in which paAIP2 was targeted to excitatory neurons using a CaMKIIα(1.3kb) promoter (paAIP2_CaMKIIα_), and 7 SST-Cre mice that received an injection of the DIO-paAIP2 vector (paAIP2_SST_). **(C)** The average population response to mismatch was stronger in control (black) than in paAIP2_CaMKIIα_ (purple) hemispheres. Shading indicates SEM across neurons. Orange shading and bar indicate duration of mismatch. Mean responses were compared across neurons in the time window marked by the black bar above the traces. Here and in subsequent panels, n.s.: p > 0.05, *: p < 0.05, **: p < 0.01, ***: p < 0.001. For all details of statistical testing, see **Table S1**. **(D)** As in **C**, but for responses to the onset of a drifting grating stimulus (see Methods). Green shading and bar indicate presence of grating stimulus. **(E)** As in **C**, but for running onset responses in closed-loop sessions. **(F)** As in **C**, but for inhibition of CaMKII in SST interneurons. **(G)** As in **D**, but for inhibition of CaMKII in SST interneurons. **(H)** As in **E**, but for inhibition of CaMKII in SST interneurons.

Given the differences between *ΔGrin1*_juv_ and the CaMKII inhibition in excitatory neurons, we compared the effect of CaMKII inhibition in inhibitory interneurons to that observed when inhibiting CaMKII in excitatory neurons. Inhibiting CaMKII in SST interneurons had an effect similar to the one we found when inhibiting CaMKII in excitatory neurons, decreasing mismatch responses and increasing visual responses (**Figures 5F-5H**). Interestingly, the running onset responses during the closed-loop session were much larger in the inhibited hemisphere (**Figure 5H**). This could be explained by an increased motor-related excitatory input, a decreased bottom-up visual inhibition, or a combination of both. Assuming SST interneurons mediate visually driven inhibition, and the establishment of this inhibition is experience dependent, we would expect that the CaMKII inhibition in SST interneurons results in decreased visually driven inhibition onto L2/3 neurons. To test this, we quantified the average correlation between neuronal activity and visual flow speed in open-loop sessions. Under normal conditions, this correlation is negative for L2/3 excitatory neurons (**Figure 6A**). The correlation became more strongly negative with the inhibition of CaMKII in excitatory neurons but became positive with the inhibition of CaMKII in SST interneurons. This positive shift in the correlation with visual flow is consistent with a decrease in visually driven inhibition by the paAIP2 inhibition of CaMKII in SST interneurons. To test whether this effect is specific to SST interneurons or simply the consequence of altering inhibition, we repeated the experiments in PV-Cre mice. Consistent with a role of PV interneurons in modulating cortical gain (Atallah et al., 2012), inhibiting CaMKII in PV interneurons resulted in a uniform increase in all response types (**Figures S3C-S3E**), but did not lead to a net positive correlation of neural activity with visual flow in excitatory neurons (**Figure 6A**). Moreover, comparing the average visual flow correlation across all manipulations, we found that only the inhibition of CaMKII in SST interneurons resulted in a net positive correlation of neural activity with visual flow in excitatory neurons. Thus, plasticity in SST interneurons is likely central to establishing normal levels of visually driven inhibition in V1.

**Figure 6.**
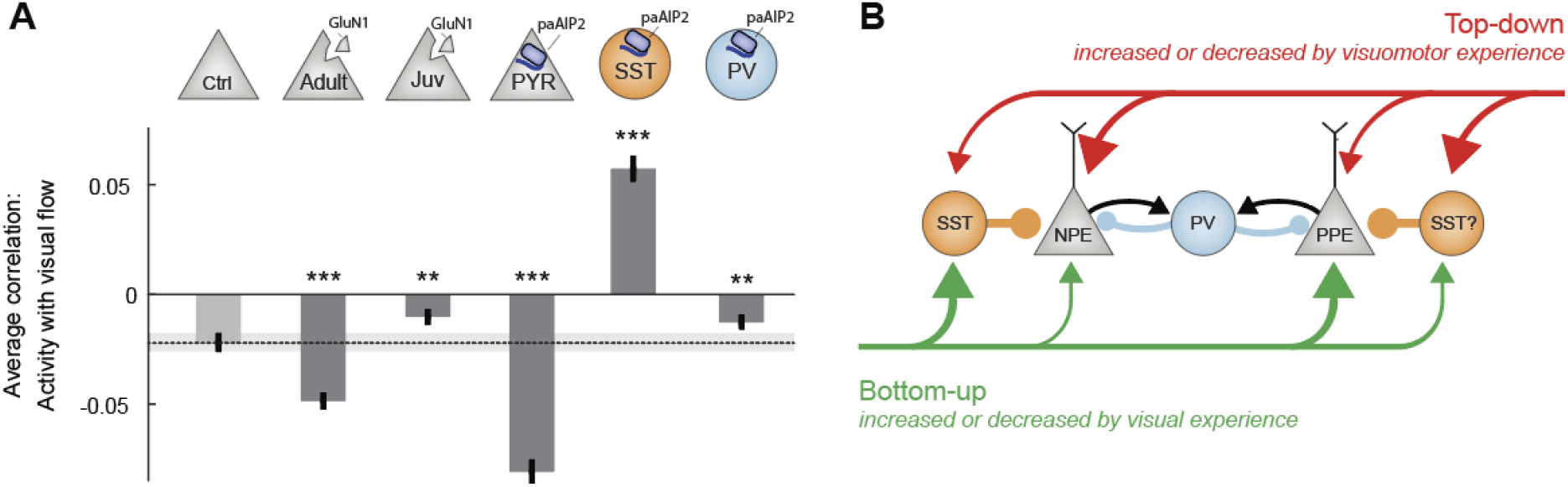
CaMKII inhibition in SST interneurons during first visuomotor experience reduced visually driven inhibition. **(A)** Mean correlation between neuronal activity and visual flow in open-loop sessions for all L2/3 excitatory neurons recorded in adult control, *ΔGrin1*_adult_, *ΔGrin1*_juv_, paAIP2_CaMKIIα_, paAIP2_SST_ and paAIP2_PV_ mice. Error bars indicate SEM across neurons. Dashed line (black) indicates mean correlation of activity and visual flow of the adult control group; gray shading indicates SEM across neurons. Comparison against adult control data: n.s.: p > 0.05, **: p < 0.01, ***: p < 0.001. For all details of statistical testing, see **Table S1**. **(B)** Through visuomotor experience, local plasticity in V1 establishes a balance between top-down and bottom-up input in L2/3 neurons (Jordan and Keller, 2020), that is thought to drive prediction error responses. In this model, we refer to neurons that receive strong bottom-up excitation and strong top-down inhibition as positive prediction error (PPE) neurons, while those with strong top-down excitation and strong bottom-up inhibition, we refer to as negative prediction error (NPE) neurons. Given that interfering with plasticity in either excitatory neurons or SST interneurons prevents normal development of visual responses in excitatory neurons, combined with the finding that visual responses in neither population of neurons depend on coupled visuomotor experience (Attinger et al., 2017), we conclude that visual experience is necessary and sufficient for shaping visual inputs onto both populations of neurons. As mismatch responses in excitatory neurons depend on visuomotor experience and are sensitive to blocking plasticity in excitatory neurons, the proper wiring of top-down input onto L2/3 excitatory neurons likely requires coupled visuomotor experience. SST interneurons likely mediate visually driven inhibition, and we speculate that they also mediate the top-down motor-related inhibition. The effect of interfering with plasticity in PV interneurons is consistent with the idea that they regulate overall gain of the circuit.

To test if normal visuomotor experience without inhibition of CaMKII would revert the changes we observed, we returned the mice to dark housing for 2 days following the first imaging session and repeated the neuronal activity measurements. At the time of the second measurement, the only visual experience without inhibition of CaMKII the mice had experienced was approximately 15 min of closed-loop visual feedback, 30 min of open-loop visual feedback, and 15 min of grating stimuli of the first imaging session. We found that after this one hour of visual experience, most of the CaMKII inhibition induced effects had either significantly reduced or reverted. For mice with inhibition of CaMKII in excitatory neurons or SST interneurons, mismatch responses in the inhibited hemisphere were larger on the 2^nd^ day of imaging than on the first day of imaging (**Figures S4A and S4D**), while grating onset responses were significantly reduced compared to the first day of imaging (**Figures S4B and S4E**). Running onset responses in the closed-loop sessions on the 2^nd^ day of imaging were decreased in the inhibited hemisphere compared to those on the first day of imaging (**Figures S4C and S4F**) and the correlation of neuronal activity with visual flow became negative in the mice that had originally received CaMKII inhibition in SST interneurons (**Figure S4G**). Thus, normal visuomotor coupling in absence of CaMKII inhibition allowed the circuit to revert towards the control state. Together, these data are consistent with the interpretation that plasticity both in the top-down input to L2/3 as well as the visually driven inhibition mediated by SST interneurons is necessary to establish the L2/3 circuit underlying the computation of visuomotor prediction errors.

## DISCUSSION

Our results demonstrate that with first visual experience in the life of a mouse, exposure to visuomotor coupling establishes a circuit in V1 capable of integrating motor and visual signals that enables visuomotor skill learning later in life. Given that the block of NMDA-receptor-dependent plasticity resulted in a reduction of responses in L2/3 neurons to mismatch and visual stimuli, we speculate that the observed impairment in visuomotor skill learning is the consequence of a reduced capacity of V1 to compute visuomotor prediction errors. Considering that L2/3 excitatory neurons balance opposing bottom-up and top-down input (Jordan and Keller, 2020), our results indicate that this balance is established by local plasticity in V1 through experience with visuomotor coupling early in life. We find that preventing this process from occurring in V1 during visuomotor development impairs the ability of mice to learn visuomotor tasks later in life. Thus, we speculate that the ability of V1 to compute visuomotor prediction errors is an essential component of the computational strategy the brain uses to guide movement by visual feedback in complex behavioral tasks. Interestingly, later in life, plasticity in V1 is no longer necessary for visuomotor skill learning, indicating that most of the learning related plasticity occurs outside of V1 or independent of NMDA receptors.

When interpreting our results, it should be kept in mind that our strategy to knock out NMDA receptors in V1 is not specific to L2/3 neurons, and we cannot be certain if the effects we observed in L2/3 neurons are the direct consequence of the NMDA receptor knockout in these neurons or a downstream consequence of an effect in another layer. It has been demonstrated, however, that a knockout of NMDA receptors in L4 neurons, the main source of bottom-up visual input to L2/3 neurons, does not alter visually evoked potentials in visual cortex, nor does it impair visual acuity of the mice, regardless of whether the knockout is congenital or postadolescent (Fong et al., 2020; Sawtell et al., 2003). Thus, we speculate that the NMDA receptor knockout effects we observe are at least in part driven by interfering with the establishment of normal input to the L2/3 neurons in V1. Another potential confound of these experiments is that we are using intracellular calcium concentration changes to measure neuronal activity, when the NMDA receptor channel is permeable to calcium and constitutes the main source of calcium in dendritic spines (Sabatini et al., 2002). However, given that we are measuring calcium signals at the soma where the main source of calcium is voltage-gated calcium channels (Grienberger and Konnerth, 2012), the direct effect of the NMDA receptor knockout on intracellular calcium is unlikely to interfere with our conclusions. Moreover, an overall reduction in calcium would influence all responses equally and would not explain why after NMDA receptor knockout, we find a strong reduction in mismatch and visual responses but only a small reduction in mean activity levels in juvenile mice (**Figure 1**), while in adult mice the converse is true (**Figure 2**).

There is a marked difference between the NMDA receptor knockout results and the CaMKII inhibition results in that the latter led to a massive increase in visual responses. There are a number of possible explanations that could account for this difference. First, despite the fact that NMDA receptors and CaMKII are closely linked in many forms of synaptic plasticity, there could be a systematic difference in the dependence of plasticity on the two molecules as a function of neuron or synapse type. Second, while the NMDA receptor knockout is permanent, we only inhibit CaMKII during the 2 hours of visuomotor exposure. Outside of this time, when the mice were housed in darkness, there could have been forms of compensatory plasticity in V1 in response to visuomotor experience driven plasticity outside of V1. Third, as the inhibition of CaMKII is driven by blue light illumination on the cortical surface, there could be a systematic difference in which synapses, or neurons, are influenced by the manipulation. The power of the light used to activate paAIP2 falls off exponentially with cortical depth with an estimated decay constant of between 35 μm and 97 μm (**Figure S2A**), consistent with previous findings (Yona et al., 2016). This, combined with the fact that CaMKII expression is higher in superficial L2/3 neurons than L4 and L5 neurons (Lein, 2007; Tighilet et al., 1998), could result in an increased effect of the CaMKII inhibition in superficial synapses. Long-range cortical input, which is thought to carry motor-related input to V1 (Leinweber et al., 2017), arrives preferentially on more superficial synapses than the bottom-up visual input (Park et al., 2019; Petreanu et al., 2009; Young et al., 2021). Thus, the differences in effect on grating responses between the NMDA receptor knockout and the CaMKII inhibition could be explained by a differential influence on top-down and bottom-up pathways. While we cannot exclude the involvement of the other potential explanations discussed above, it is not immediately clear why they would result in a differential effect with regards to positive and negative prediction errors. Thus, we speculate that the CaMKII inhibition results in a differential impairment of plasticity in top-down and bottom-up pathways.

It is important to note that the within animal control suffers from the confound that the two hemispheres are directly connected. For instance, the fact that visual responses are also massively increased in the control hemisphere of mice in which we inhibited CaMKII in excitatory neurons relative to the level of responses one would expect normally (e.g., compare **Figure 5D** with **Figure 2B**, or (Attinger et al., 2017)), is likely caused by this direct interaction. A similar problem befalls our experiments using the NMDA receptor knockout. However, given that the effect sizes were considerably smaller in those experiments, crosstalk effects are likely also less salient.

Lastly, given that little is known about the role of CaMKII in the plasticity in interneurons, it was not *a priori* clear that blocking CaMKII in SST or PV interneurons during visuomotor development would have a measurable influence on L2/3 excitatory neuron responses. While CaMKIIα is mainly expressed in excitatory neurons in cortex, CaMKIIβ is found in both excitatory and inhibitory neurons (Nicole and Pacary, 2020). Given that paAIP2 is designed based on a sequence of the autoinhibitory domain of CaMKII (Hanson et al., 1989) that is highly conserved across isoforms (Tobimatsu and Fujisawa, 1989), and inhibits CaMKII at the kinase domain (Murakoshi et al., 2017), which is also highly conserved across isoforms (Tobimatsu and Fujisawa, 1989), paAIP2 inhibition is likely independent of CaMKII isoform. Thus, our results would be consistent with the interpretation that SST and PV interneurons exhibit CaMKIIβ-dependent forms of plasticity necessary for the establishment of normal visuomotor integration in V1. Supporting this interpretation is the fact that inhibiting CaMKII in SST interneurons has an effect on net visual drive opposite to that of the same inhibition in excitatory neurons (**Figure 6A**). Consistent with the previous finding that SST activity is critical for the computation of visuomotor mismatch responses (Attinger et al., 2017), the role of SST interneurons appears to be critical to establishing a balance between top-down and bottom-up input in L2/3 neurons in V1. This is in line with the findings that the activity of SST interneurons is modulated by locomotion only in the presence of visual input (Pakan et al., 2016), and that in excitatory neurons, inputs from both excitatory neurons and SST interneurons, but not PV or vasoactive intestinal peptide (VIP)-expressing interneurons, exhibit NMDA receptor-dependent plasticity (Chiu et al., 2018). Thus, we postulate that visuomotor experience establishes a balance in individual L2/3 neurons, either between top-down excitatory input and visually driven inhibition mediated by SST interneurons, or top-down inhibition - possibly also mediated by SST interneurons - and visually driven excitation (**Figure 6B**).

## ABBREVIATIONS

CaMKII: Calcium/calmodulin-dependent protein kinase II
Grin1: Glutamate receptor ionotropic NMDA subunit 1
L2/3: Layer 2/3 of cortex
L4: Layer 4 of cortex
L5: Layer 5 of cortex
NMDA: N-Methyl-D-aspartate
paAIP2: photoactivatable autocamtide inhibitory peptide 2 (an inhibitor of CaMKII)
PV: Parvalbumin
SEM: Standard error of the mean
SST: Somatostatin
V1: Primary visual cortex

## SUPPLEMENTARY FIGURES

**Figure S1.**
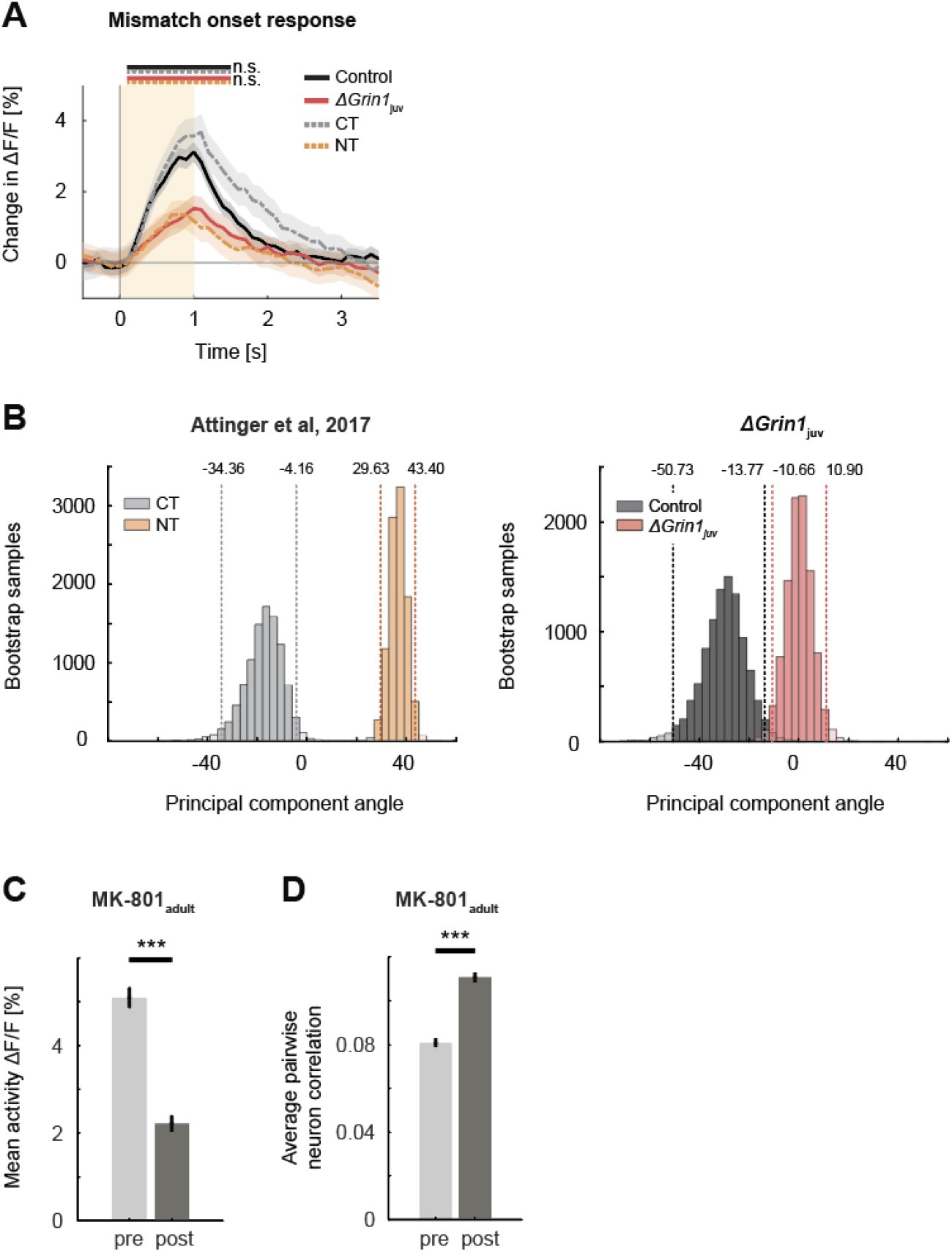
The effect of the NMDA receptor knockout was comparable to the lack of experience with visuomotor coupling and systemic block of NMDA receptors. Related to Figure 1. **(A)** Mean population response to mismatch in *ΔGrin1*_juv_ (red) hemisphere, the control hemisphere (black), coupled trained controls (CT, dashed gray), and mice raised without visuomotor coupling (non-coupled trained (NT), orange). Responses in *ΔGrin1*_juv_ are similar to those in NT mice, and those in the control hemisphere were similar to those in CT mice. Note, the data shown in this figure includes all data, while the data shown in **Figure 1E** includes only those data from recording sites for which we also had sufficient grating and running onset data (see Methods). Here and in subsequent panels, n.s.: p > 0.05, *: p < 0.05, **: p < 0.01, ***: p < 0.001. For all details of statistical testing, see **Table S1**. **(B)** Bootstrap distribution of principal component angles from correlational analysis of neuronal activity with running speed and visual flow. Data shown are from coupled trained (CT, dashed gray) and non-coupled trained (NT, dashed orange) mice from (Attinger et al., 2017), and for *ΔGrin1*_juv_ (solid red) and control hemisphere (solid black) data. **(C)** Mean activity of L2/3 neurons in V1 before (pre, light gray) and 1 hour after (post, dark gray) intra-peritoneal injection of the NMDA receptor antagonist MK-801 (0.1 mg/kg). Error bars indicate SEM across neurons. **(D)** Average pairwise correlation of neuronal activity was higher 1 hour after MK-801 injection (post, dark gray) compared to pre injection. Error bars indicate SEM across neurons.

**Figure S2.**
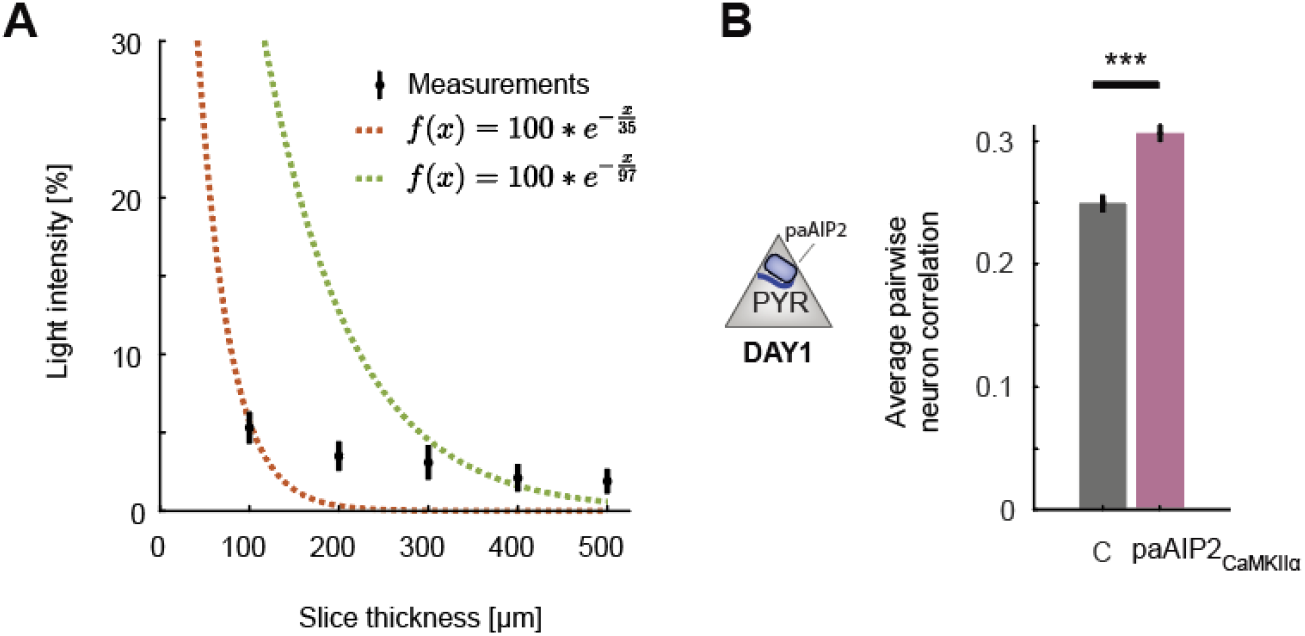
Additional data of CaMKII inhibition in excitatory or SST inhibitory neurons. Related to Figure 5. **(A)** Percentage of blue light (473 nm) power transmitted through acute slices of cortical tissue of varying thickness. Shown in black are mean and standard deviation over 6 measurements. The dashed red line is a least squares exponential fit with a decay constant of 37 μm, and the green line is the transform of a least squares linear fit to the log-transformed data with a decay constant of 97 μm. Note, the data are not well fit by an exponential decay likely as a result of the point illumination. See (Yona et al., 2016) for detailed modelling of power decay. **(B)** Average pairwise correlation of neuronal activity is higher in CaMKII inhibited excitatory neurons (purple), compared to that in the uninhibited control hemisphere (black). Error bars indicated SEM. Here and in subsequent panels, n.s.: p > 0.05, *: p < 0.05, **: p < 0.01, ***: p < 0.001. For all details of statistical testing, see **Table S1**.

**Figure S3.**
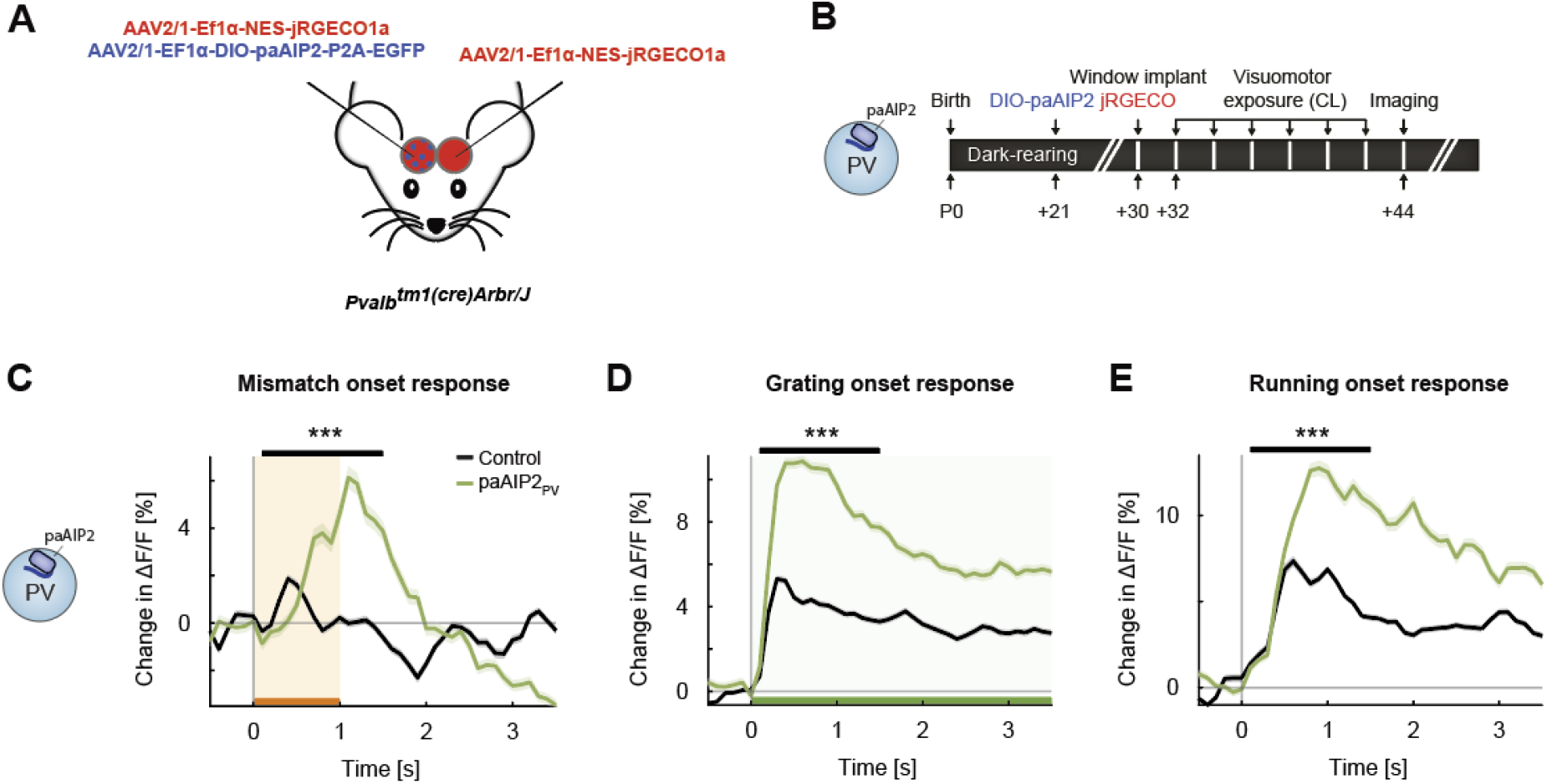
Inhibiting CaMKII in PV interneurons resulted in an overall increase in onset responses in L2/3 excitatory neurons. Related to Figure 5. **(A)** We unilaterally injected an eGFP-tagged paAIP2 or DIO-paAIP2 expressing virus and a calcium indicator (jRGECO1a) bilaterally in V1. **(B)** 6 PV-Cre mice were dark reared from birth. AAV injections occurred at postnatal day 21 (DIO-paAIP2) and P30 (jRGECO1a). Imaging window implantation occurred on P30. Mice had 6 closed-loop training sessions (visuomotor exposure) during which we illuminated cortex bilaterally with blue light (473 nm) to inhibit CaMKII. **(C)** The average population response to mismatch was stronger in the paAIP2_PV_ (green) than in the control (black) hemispheres. Orange shading and bar indicate duration of mismatch. Shading indicates SEM. Mean responses are compared across neurons in the time window indicated by the black bar above the traces. Here and in subsequent panels, n.s.: p > 0.05, *: p < 0.05, **: p < 0.01, ***: p < 0.001. For all details of statistical testing, see **Table S1**. **(D)** As in **C**, but for responses to the onset of a drifting grating stimulus (see Methods). Green shading and bar indicate presence of grating stimulus. **(E)** As in **C**, but for running onset responses in closed-loop sessions.

**Figure S4.**
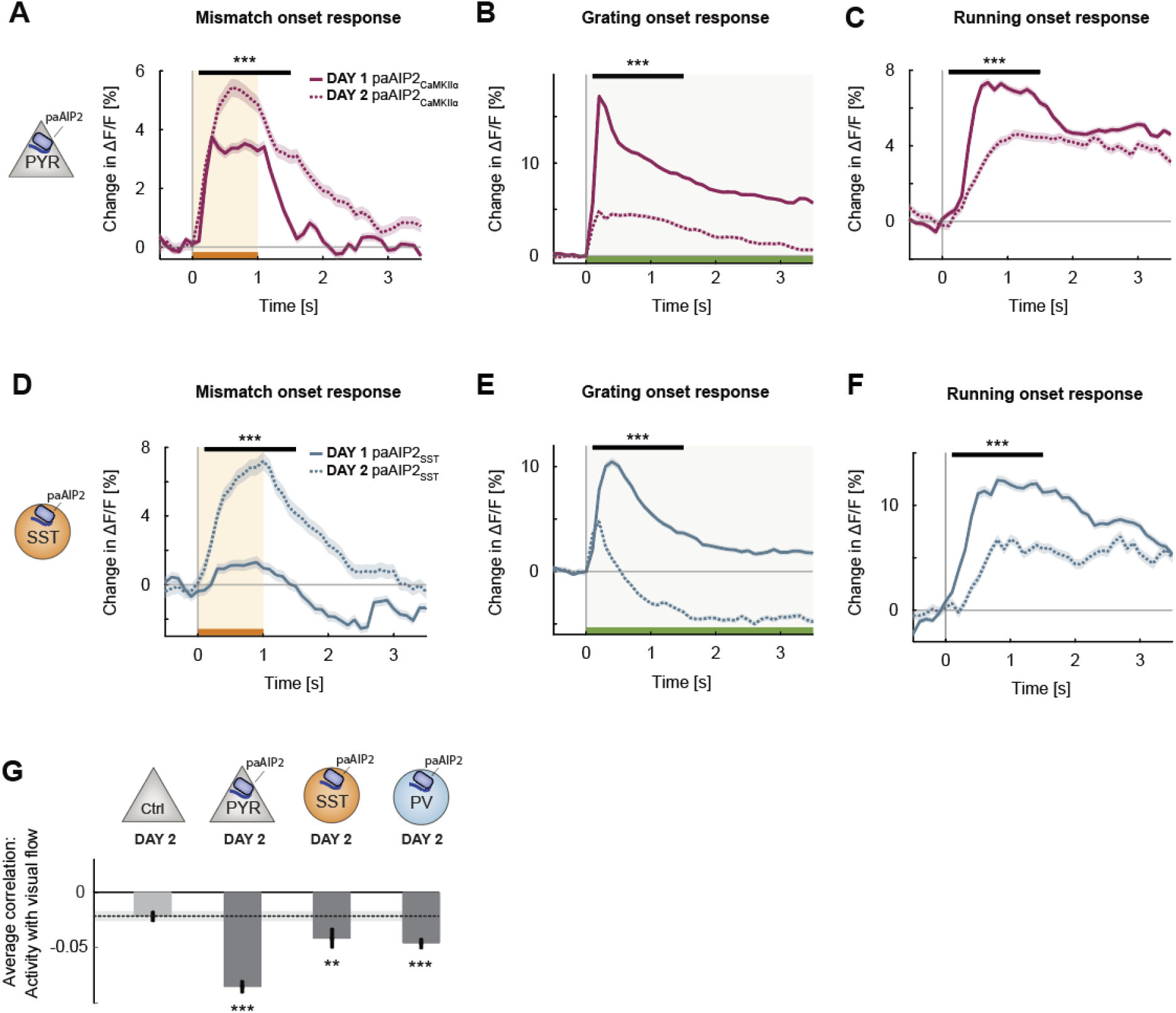
Changes induced by CaMKII inhibition quickly reverted with exposure to normal visuomotor coupling. Related to Figure 5. **(A)** The average population response to mismatch on day 2 of imaging (dashed) and on day 1 of imaging (solid). Shading indicates SEM. Orange shading and bar indicate duration of mismatch. Mean responses are compared across neurons in the time window indicated by the black bar above the traces. Here and in subsequent panels, n.s.: p > 0.05, *: p < 0.05, **: p < 0.01, ***: p < 0.001. For all details of statistical testing, see **Table S1**. **(B)** As in **A**, but for responses to the onset of a drifting grating stimulus (see Methods). Green shading and bar indicate presence of grating stimulus. **(C)** As in **A**, but for running onset responses in closed-loop sessions. **(D)** As in **A**, but for inhibition of CaMKII in SST interneurons using paAIP2. **(E)** As in **B**, but for inhibition of CaMKII in SST interneurons using paAIP2. **(F)** As in **C**, but for inhibition of CaMKII in SST interneurons using paAIP2. **(G)** Mean correlation between neuronal activity and visual flow in open-loop sessions for all L2/3 neurons recorded in the paAIP2 inhibited hemispheres of paAIP2_CaMKIIα_, paAIP2_SST_, and paAIP2_PV_ mice on day 2, compared to the responses in the adult control group. Error bars indicate SEM across neurons. Dashed line (black) indicates mean correlation of activity and visual flow of the adult control group; gray shading indicates SEM across neurons. Comparison against normally reared, adult control data: n.s.: p > 0.05, **: p < 0.01, ***: p < 0.001.

## METHODS

**Key Resources Table.**

| REAGENT or RESOURCE | Source | Identifier |
| --- | --- | --- |
| **Bacterial and Virus Strains** |  |  |
| AAV2/1-EF1α-GCaMP6f-WPRE (6.0 10<sup>11</sup> - 8.0 10<sup>12</sup> GC/ml) | FMI vector core | vector.fmi.ch |
| AAV2/1-EF1α-Cre-t2a-mcherry-WPRE (3.2 10<sup>11</sup> – 1.2 10<sup>13</sup> GC/ml) | FMI vector core | vector.fmi.ch |
| AAV2/1-EF1α-Cre-WPRE (2.810<sup>10</sup> GC/ml) | FMI vector core | vector.fmi.ch |
| AAV2/1-EF1α-NES-jRGECO1a-WPRE (4.8 10<sup>13</sup> GC/ml) | FMI vector core | vector.fmi.ch |
| AAV2/1-CaMKIIα(1.3kb)-mEGFP-P2A-paAIP2 (1.80 10<sup>13</sup> GC/ml) | FMI vector core | vector.fmi.ch |
| AAV2/1-EF1α-DIO-mEGFP-P2A-paAIP2-WPRE (1.2 10<sup>13</sup> GC/ml) | FMI vector core | vector.fmi.ch |
| **Chemicals, Peptides, and Recombinant Proteins** |  |  |
| Fentanyl citrate | Actavis | CAS 990-73-8 |
| Midazolam (Dormicum) | Roche | CAS 59467-96-8 |
| Medetomidine (Dormitor) | Orion Pharma | CAS 86347-14-0 |
| Ropivacaine | Presenius Kabi | CAS 132112-35-7 |
| Lidocaine | Bichsel | CAS 137-58-6 |
| Buprenorphine | Reckitt Benckiser Healthcare | CAS 52485-79-7 |
| Ophthalmic gel (Humigel) | Virbac | N/A |
| Flumazenil (Anexate) | Roche | CAS 78755-81-4 |
| Atipamezole (Antisedan) | Orion Pharma | CAS 104054-27-5 |
| N-Butyl-2-cyanoacrylate (Histoacryl) | Braun | CAS 6606-65-1 |
| Dental cement (Paladur) | Heraeus Kulzer | CAS 9066-86-8 |
| MK-801 | Sigma | CAS 77086-22-7 |
| **Deposited data** |  |  |
| All data and code used to generate manuscript figures | This paper | data.fmi.ch |
| **Experimental Models: Organisms/Strains** |  |  |
| *Mus musculus*: C57BL/6J | Charles River | N/A |
| *Mus musculus*: *Grin1*<sup>tm2Stl/J</sup> | Jackson laboratories | Cat#005246 |
| *Mus musculus*: *Pvalb*<sup>tm1(cre)Arbr</sup> | Jackson laboratories | Cat#008069 |
| *Mus musculus*: *Sst*<sup>tm2.1(cre)Zjh</sup> | Jackson laboratories | Cat#018973 |
| **Software and algorithms** |  |  |
| MATLAB (2020B) | The MathWorks | RRID: SCR_001622 |
| LabView | National Instruments | RRID:SCR_014325 |
| Two-photon acquisition software | Keller laboratory | sourceforge.net/p/iris-scanning/ |
| Image data processing software | Keller laboratory | sourceforge.net/p/iris-scanning/calliope |
| Python | python.org | RRID:SCR_008394 |
| Panda3D | panda3d.org | N/A |
| **Other** |  |  |
| mRNA probe Mm-Grin1-O1 (probe region 2892 - 4127) | ACD bio | Cat#473079 |
| Virtual reality and two-photon setup | (Leinweber et al., 2014, 2017) | DOI: 10.3791/50885, 10.1016/j.neuron.2017.08.036 |
| Opto-stimulation laser (OBIS 473 nm LX) | Coherent | Cat#1187194 |
| Titanium headplate | FMI/ETHZ workshop | N/A |
| Dental drill | Meisinger | N/A |

### Animals and surgery

All animal procedures were approved by and carried out in accordance with guidelines of the Veterinary Department of the Canton Basel-Stadt, Switzerland. For all surgical procedures, mice were anesthetized with a mixture of Fentanyl (0.05 mg/kg; Actavis), Midazolam (5.0 mg/kg; Dormicum, Roche) and Medetomidine (0.5 mg/kg; Domitor, Orion). Analgesics were applied perioperatively (2% Lidocaine gel, Meloxicam 5mg/kg) and post-operatively (Buprenorphine 0.1 mg/kg, Metacam 5 mg/kg). Eyes were covered with ophthalmic gel (Virbac Schweiz AG). At postnatal day P21, we injected approximately 100 nl of AAV2/1-Ef1α-Cre-T2A-mCherry vector at a titer of between 3.2 10^11^ and 1.2 10^13^ GC/ml, or AAV2/1-EF1α-Cre-WPRE vector at a titer of 2.8 10^10^ GC/ml (**Figures 1–4**); AAV2/1-CaMKIIα(1.3kb)-mEGFP-P2A-paAIP2 vector at a titer of 1.8 10^13^ GC/ml, or AAV2/1-EF1α-DIO-mEGFP-P2A-paAIP2-WPRE vector at a titer of 1.2 10^13^ GC/ml (**Figure 5**) through a small hole in the skull over the right hemisphere at 2.4 mm directly lateral from lambda.

For window implantations at P30, we performed a cranial window surgery by implanting a circular 4 mm glass coverslip bilaterally, following injections of approximately 200 nl of AAV vectors (AAV2/1-EF1α-GCaMP6f-WPRE or AAV2/1-EF1α-NES-jRGECO1a-WPRE) into V1, 2.5 mm lateral from lambda.

### Virtual reality environment and virtual navigation task

In all experiments involving the virtual reality system, mice were head-fixed and mounted on a spherical treadmill, as described previously (Leinweber et al., 2014). In brief, mice were free to run on an air supported polystyrene ball. Ball rotation controlled movement in a virtual reality environment displayed on a toroidal screen surrounding the mouse and covered approximately 240 degrees horizontally and 100 degrees vertically of visual space, from the point of view of the mouse.

First visual and visuomotor exposure of the mice occurred in this virtual reality environment in the 12 days prior to imaging experiments. Mice were trained for 2 hours every other day (for a total of 6 sessions) with closed-loop feedback between forward locomotion and backward visual flow in a virtual corridor with walls textured with vertical sinusoidal gratings (Attinger et al., 2017). All two-photon imaging experiments were also performed on the same virtual reality setup, and unless otherwise noted, data were acquired in sessions of 5-15 minutes duration in the following sequence: Closed-loop, open-loop, dark, grating. In closed-loop sessions, running was coupled to movement in the same virtual environment used during visuomotor exposure. In open-loop sessions, the self-generated visual flow from the preceding closed-loop session was replayed. In grating sessions, drifting grating stimuli of different directions (0, 45, 90, 270 degrees, moving in either direction) were presented in random sequences. Each grating lasted between 3 s to 8 s with an inter-trial interval (gray screen) of between 2 s and 6 s. For all experiments, rotation of the ball was restricted to the transverse axis to allow only forward and backward movement in the virtual reality environment. Mice were free to run in all experiments and did so spontaneously.

For the virtual navigation experiments (**Figure 4**), rotation of the ball was not restricted, and mice could control forward and backward motion, as well as rotation in the virtual environment. To incentivize mice to engage in the visuomotor skill learning task, they were water-restricted with access to 1 mL water daily 3 days before the start of the behavioral experiments. Care was taken to prevent a drop in body weight to below 80% throughout training. During the experiment, mice could obtain water rewards by reaching the end of the virtual corridor, after which they were presented with a 5 second gray screen and teleported to the beginning of the corridor again. Task difficulty was increased with increasing performance of the mice by expanding the length of the virtual corridor to keep the rate of water rewards roughly constant. At the beginning of training the length-to-width ratio of the corridor was 5. Every 4 trials, the length of the corridor would be updated by a factor between 1 (no change) and 1.5 (50% increase in length), where the factor was determined as 20 s divided by the mean duration of those 4 trails. Maximum corridor length was restricted to 400% of the length on the first day. Visual offset perturbations were introduced once per trial, presented at a random position within 20% and 80% of the total corridor length and consisted of 30° heading offsets introduced randomly, either to the left or to the right. The task performance index (PI) was calculated as follows:

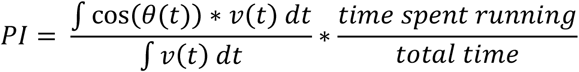

Where θ(t) is the direction of running relative to the target, and v(t) is the running speed of the mouse. The intuition behind this index is to quantify performance as the fraction of distance traveled in the direction of the target, normalized by the total distance traveled. The second factor is added to reduce variability driven by a short time spent running, as is typical in early training sessions.

### Two-photon calcium imaging

Two-photon imaging of L2/3 neurons in V1 was performed as described previously (Leinweber et al., 2014, 2017). In brief, two-photon imaging was done using a modified Thorlabs Bergamo I or II microscope. Excitation light source was a tunable, femtosecond-pulsed laser (Insight, Spectra Physics, tuned to 910 nm or 980 nm for GCaMP6f excitation, and 1030 nm for jRGECO1a excitation). The scan head was based on an 8 kHz or 12 kHz resonant scanner (Cambridge Technology). We used a piezo electric linear actuator (P-726, Physik Instrumente) to sequentially image 4 z-planes (approximately 40 μm apart) by moving a 16x, 0.8 NA objective (Nikon N16XLWD-PF). Emission light was band-pass filtered using a 525/50 nm or a 607/70 nm filter (Semrock), detected by a photomultiplier tube (PMT, H7422P, Hamamatsu), amplified (DHPCA-100, Femto), digitized at 800 MHz (NI5772, National Instruments) and band-pass filtered at 80 MHz using digital Fourier transform on a field-programmable gate array (NI5772, National Instruments, loaded with custom-designed logic). Images were acquired at 750 by 400 pixels using custom-written LabView software (available on a public SourceForge repository, see Key Resources table), with 10 or 15 frames per z-plane per second and a field of view of approximately 375 μm by 300 μm. Whenever possible, imaging was performed in both control and intervention hemisphere in each mouse. In a subset of mice, (see **Table S2**) imaging was only possible in one hemisphere as imaging quality did not meet our minimum quality standards in the other (clear image visible in single frame at less than 60 mW total laser power).

### Conditional *Grin1* knockout, histology, and pharmacological NMDA receptor inhibition

All *ΔGrin1* knockout experiments were performed using the *Grin1^tm2Stl^* (also known as fNR1 or NR1^flox^) mouse line (Tsien et al., 1996), which has a pair of loxP sites flanking the transmembrane domain and C-terminal region of the *Grin1* gene that codes for GluN1 (also referred to as NR1), a subunit essential to the NMDA receptor (Monyer et al., 1994). We confirmed the knockout using mRNA in situ hybridization (RNAscope, Ventana) in a separate cohort of 3 mice, 14 days after injection of an AAV vector expressing Cre recombinase in both juvenile and adult mice (*ΔGrin1*_juv, adult_). We followed a standardized formaldehyde-fixed paraffin-embedded protocol. In brief, mice were transcardially perfused with phosphate buffered saline (PBS), followed by perfusion with a solution of 4% paraformaldehyde (PFA) in PBS. Brains were isolated, post-fixed overnight in 4% PFA, paraffinized for 24 h, and cut at 5 μm using a microtome (ThermoFisher). Slices were stained using hematoxylin to mark cell bodies, and Mm-*Grin1*-O1 (#473079, target region 2892 - 4127, ACDBio) to label *Grin1* mRNA. To ease identification of the knockout area in two-photon microscopy, a vector co-expressing a red fluorophore (mCherry) and Cre was used to induce the *Grin1* knockout in most experiments. Due to a shortage of the correct vector, a subset (6 of 14) of the *ΔGrin1*_adult_ experiments were performed without the mCherry fluorophore. For pharmacological NMDA receptor inhibition experiments (**Figures S1C-D**), adult wild-type mice were injected with 0.1 mg/kg MK-801 intraperitoneally and neuronal activity was recorded before and after injection.

### Optogenetic activation of paAIP2 and laser attenuation measurements

We used a photoactivatable autocamtide inhibitory peptide 2 (paAIP2) (Murakoshi et al., 2017) to inhibit calcium/calmodulin-dependent kinase II (CaMKII) for the entire duration of the visuomotor exposure in the virtual reality environment. We directed a blue laser (OBIS 473 nm LX 75 mW, Coherent) onto V1 in both hemispheres using a galvo-galvo system and a set of mirrors and lenses (GVSM002-EC/M, Thorlabs). Beam diameter on the cortical surface was 3 mm. Light was triggered at 0.2 Hz with a duty cycle of 20% (1 s on, 4 s off). During illumination periods, we alternated between the two hemispheres at 50 Hz. Peak laser power was 20 mW, which resulted in a time-averaged power density at the cortical surface of 0.28 mW/mm^2^. To measure the laser attenuation through tissue (**Figure S2A**), we prepared slices of fresh brain tissue of 100 μm, 200 μm, 300 μm, 400 μm and 500 μm thickness. We then illuminated slices with the blue laser used for optogenetic inhibition of CaMKII set to 20 mW power and measured the fraction of power transmitted through each slice using a power meter (PM100D, Thorlabs).

### Data analysis

Calcium imaging data were processed as described previously (Keller et al., 2012). In brief, raw images were full-frame registered to correct for lateral brain motion. Neurons were selected manually based on mean and maximum fluorescence images. Average fluorescence per neurons over time was corrected for slow fluorescence drift using an 8^th^ percentile filter and a 100 s window (Dombeck et al., 2007) and divided by the median value over the entire trace to calculate ΔF/F.

Data analysis was performed with custom analysis scripts written in MATLAB 2020b (MathWorks). For all population onset responses, data were first averaged over onsets for each neuron and then averaged over neurons. Unless stated otherwise, shading and error bars indicate the standard error of the mean (SEM) across neurons. We did not test for normality of distributions. For analysis of onset responses (**Figures 1E-1G**, **Figures 2A-2C**, **Figures 5C-5H**, **Figures S4A-S4F**), recording sites with less than 3 running or mismatch onsets in a particular session were excluded from analysis (e.g., if a mouse ran without stopping for the entire closed-loop session, there were no running onsets to analyze). We also excluded 2 sessions in which the mouse did not run without prompting by the experimenter. In total, we excluded 10 of 128 sessions. In 31 of 118 of the remaining sessions, we did not record grating responses. To calculate stimulus induced changes in ΔF/F, we used a baseline subtraction window of −300 ms to 0 ms, and a response window of +100 ms to +1500 ms relative to stimulus onset. To determine running onsets, we used a threshold of 10^−2^ cm/s. For analysis of average activity levels in closed-loop sessions (**Figure 1H**, **Figure 2D**, **Figure S1C**), we calculated the average neuronal activity (ΔF/F [%]) over time and over neurons. To calculate average pairwise correlation between neurons in closed-loop sessions (**Figure 1J**, **Figure 2F**, **Figure S1D**, **Figures S2B**), we calculated the mean correlation of each neuron with all other neurons. To calculate the first principal component (**Figure 1I**, **Figure 2E**, **Figure S1B**), we calculated the eigenvectors of the covariance matrix of the mean-subtracted visual flow and running correlations with neuronal activity. The principal component angle was defined as the angle between the first principal component and the y axis.

**Table S1.** Statistics. All values are rounded to two significant decimals, except values smaller than 10^−5^. The tests used, were two-sample independent t-test (t-test 2), one sample t-test (t-test 1), or a bootstrap estimate of the 95% confidence interval (CI).

| Reference | Description | Test | N<sub>1</sub> | N<sub>2</sub> | Unit | P-value/CI |
| --- | --- | --- | --- | --- | --- | --- |
| **Figure 1E-G** | Control (N<sub>1</sub>) vs $\Delta Grin1_{juv}$ (N<sub>2</sub>) | | | | | |
|  | Mismatch response | t-test 2 | 794 | 551 | neurons | 0.000082 |
|  | Drifting grating response | t-test 2 | 794 | 551 | neurons | 0.030 |
|  | Running onset response (closed-loop) | t-test 2 | 794 | 551 | neurons | 0.054 |
| **Figure 1H** | Control (N<sub>1</sub>) vs $\Delta Grin1_{juv}$ (N<sub>2</sub>) | t-test 2 | 2625 | 1986 | neurons | 0.81 |
| **Figure 1J** | Control (N<sub>1</sub>) vs $\Delta Grin1_{juv}$ (N<sub>2</sub>) | t-test 2 | 2625 | 1986 | neurons | <10<sup>-5</sup> |
| **Figure 2A-C** | $\Delta Grin1_{adult}$ (N<sub>2</sub>) vs control (N<sub>1</sub>) | | | | | |
|  | Mismatch response | t-test 2 | 912 | 1184 | neurons | 0.45 |
|  | Drifting grating response | t-test 2 | 912 | 1184 | neurons | 0.57 |
|  | Running onset response (closed-loop) | t-test 2 | 912 | 1184 | neurons | 0.45 |
| **Figure 2D** | Control (N<sub>1</sub>) vs $\Delta Grin1_{adult}$ (N<sub>2</sub>) | t-test 2 | 1281 | 1547 | neurons | <10<sup>-5</sup> |
| **Figure 2F** | Control (N<sub>1</sub>) vs $\Delta Grin1_{adult}$ (N<sub>2</sub>) | t-test 2 | 1281 | 1547 | neurons | <10<sup>-5</sup> |
| **Figure 3C** | Control<sub>adult</sub> (N<sub>1</sub>) vs |  |  |  |  |  |
| | $\Delta Grin1_{juv}$ control (N<sub>2</sub>) | t-test 2 | 869 | 794 | neurons | 0.15 |
| | $\Delta Grin1_{juv}$ (N<sub>2</sub>) | t-test 2 | 869 | 551 | neurons | 0.000038 |
| | $\Delta Grin1_{adult}$ control (N<sub>2</sub>) | t-test 2 | 869 | 912 | neurons | 0.37 |
| | $\Delta Grin1_{adult}$ (N<sub>2</sub>) | t-test 2 | 869 | 986 | neurons | 0.24 |
| **Figure 4C (top)** | $\Delta Grin1_{juv}$ average performance index on day 1-2 vs 6-7 | t-test 1 | 6 | 6 | mice | 0.22 |
| | $\Delta Grin1_{adult}$ average performance index on day 1-2 vs 6-7 | t-test 1 | 13 | 13 | mice | 0.00015 |
|  | Control<sub>juv</sub> average performance index on day 1-2 vs 6-7 | t-test 1 | 6 | 6 | mice | 0.0037 |
| **Figure 4C (right)** | Average performance index on day 6-7 |  |  |  |  |  |
| | Control<sub>juv</sub> (N<sub>1</sub>) vs $\Delta Grin1_{juv}$ (N<sub>2</sub>) | t-test 2 | 6 | 6 | mice | 0.0013 |
| | Control<sub>juv</sub> (N<sub>1</sub>) vs $\Delta Grin1_{adult}$ (N<sub>2</sub>) | t-test 2 | 6 | 13 | mice | 0.016 |
| | $\Delta Grin1_{juv}$ (N<sub>1</sub>) vs $\Delta Grin1_{adult}$ (N<sub>2</sub>) | t-test 2 | 6 | 13 | mice | 0.39 |
| **Figure 4E** | Control<sub>juv</sub> (N<sub>1</sub>) vs $\Delta Grin1_{juv}$ (N<sub>2</sub>) | t-test 2 | 6 | 6 | mice | 0.027 |
| | Control<sub>juv</sub> (N<sub>1</sub>) vs $\Delta Grin1_{adult}$ (N<sub>2</sub>) | t-test 2 | 6 | 13 | mice | 0.17 |
| | $\Delta Grin1_{juv}$ (N<sub>1</sub>) vs $\Delta Grin1_{adult}$ (N<sub>2</sub>) | t-test 2 | 6 | 13 | mice | 0.028 |
|  | Control<sub>juv</sub> (N<sub>1</sub>) vs 0 | t-test 1 | 6 | N/A | mice | 0.0057 |
| | $\Delta Grin1_{juv}$ (N<sub>1</sub>) vs 0 | t-test 1 | 6 | N/A | mice | 0.91 |
| | $\Delta Grin1_{adult}$ (N<sub>1</sub>) vs 0 | t-test 1 | 13 | N/A | mice | 0.0019 |
| **Figure 5C-E** | Control (N<sub>1</sub>) vs paAIP2<sub>CaMKIIα</sub> (N<sub>2</sub>) |  |  |  |  |  |
|  | Mismatch response | t-test 2 | 781 | 928 | neurons | <10<sup>-5</sup> |
|  | Drifting grating response | t-test 2 | 781 | 928 | neurons | <10<sup>-5</sup> |
|  | Running onset response (closed-loop) | t-test 2 | 781 | 928 | neurons | 0.0013 |
| **Figure 5F-H** | Control (N<sub>1</sub>) vs paAIP2<sub>SST</sub> (N<sub>2</sub>) |  |  |  |  |  |
|  | Mismatch response | t-test 2 | 1277 | 807 | neurons | <10<sup>-5</sup> |
|  | Drifting grating response | t-test 2 | 1277 | 807 | neurons | <10<sup>-5</sup> |
|  | Running onset response (closed-loop) | t-test 2 | 1277 | 807 | neurons | <10<sup>-5</sup> |
| **Figure 6A** | Control<sub>adult</sub> (N<sub>1</sub>) vs |  |  |  |  |  |
| | $\Delta Grin1_{adult}$ (N<sub>2</sub>) | t-test 2 | 1000 | 1516 | neurons | <10<sup>-5</sup> |
| | $\Delta Grin1_{juv}$ (N<sub>2</sub>) | t-test 2 | 1000 | 1547 | neurons | 0.0032 |
|  | paAIP2<sub>CaMKIIα</sub> (N<sub>2</sub>) | t-test 2 | 1000 | 928 | neurons | <10<sup>-5</sup> |
|  | paAIP2<sub>SST</sub> (N<sub>2</sub>) | t-test 2 | 1000 | 1022 | neurons | <10<sup>-5</sup> |
|  | vs paAIP2<sub>PV</sub> (N<sub>2</sub>) | t-test 2 | 1000 | 1705 | neurons | 0.013 |
| **Figure S1A** | CT (N<sub>1</sub>) vs $\Delta Grin1_{juv}$ control (N<sub>2</sub>) | t-test 2 | 2259 | 2080 | neurons | 0.14 |
| | NT (N<sub>1</sub>) vs $\Delta Grin1_{juv}$ (N<sub>2</sub>) | t-test 2 | 2103 | 1516 | neurons | 0.57 |
| **Figure S1B (left)** | CT | CI | 10<sup>4</sup> | N/A | PCA angles | [-34.36, -4.16] |
|  | NT | CI | 10<sup>4</sup> | N/A | PCA angles | [ 29.63, 43.40] |
| **Figure S1B (right)** | $\Delta Grin1_{juv}$ control | CI | 10<sup>4</sup> | N/A | PCA angles | [-50.73, -13.77] |
| | $\Delta Grin1_{juv}$ | CI | 10<sup>4</sup> | N/A | PCA angles | [-10.66, 10.90] |
| **Figure S1C** | Pre (N<sub>1</sub>) vs post (N<sub>2</sub>) MK-801 | t-test 2 | 2443 | 2443 | neurons | <10<sup>-5</sup> |
| **Figure S1D** | Pre (N<sub>1</sub>) vs post (N<sub>2</sub>) MK-801 | t-test 2 | 2443 | 2443 | neurons | <10<sup>-5</sup> |
| **Figure S2B** | Control (N<sub>1</sub>) vs paAIP2<sub>CaMKIIα</sub> (N<sub>2</sub>) | t-test 2 | 781 | 928 | neurons | <10<sup>-5</sup> |
| **Figure S3C-E** | Control (N<sub>1</sub>) vs paAIP2<sub>PV</sub> (N<sub>2</sub>) |  |  |  |  |  |
|  | Mismatch response | t-test 2 | 1256 | 919 | neurons | <10<sup>-5</sup> |
|  | Drifting grating response | t-test 2 | 1256 | 919 | neurons | <10<sup>-5</sup> |
|  | Running onset response (closed-loop) | t-test 2 | 1256 | 919 | neurons | <10<sup>-5</sup> |
| **Figure S4A-C** | paAIP2<sub>CaMKIIα</sub> day 1 (N<sub>1</sub>) vs paAIP2<sub>CaMKIIα</sub> day 2 (N<sub>2</sub>) |  |  |  |  |  |
|  | Mismatch response | t-test 2 | 928 | 926 | neurons | <10<sup>-5</sup> |
|  | Drifting grating response | t-test 2 | 928 | 926 | neurons | <10<sup>-5</sup> |
|  | Running onset response (closed-loop) | t-test 2 | 928 | 926 | neurons | <10<sup>-5</sup> |
| **Figure S4D-F** | paAIP2<sub>SST</sub> day 1 (N<sub>1</sub>) vs paAIP2<sub>CaMKIIα</sub> day 2 (N<sub>2</sub>) |  |  |  |  |  |
|  | Mismatch response | t-test 2 | 807 | 397 | neurons | <10<sup>-5</sup> |
|  | Drifting grating response | t-test 2 | 807 | 397 | neurons | <10<sup>-5</sup> |
|  | Running onset response (closed-loop) | t-test 2 | 807 | 397 | neurons | <10<sup>-5</sup> |
| **Figure S4G** | Control<sub>adult</sub> (N<sub>1</sub>) vs |  |  |  |  |  |
|  | paAIP2<sub>CaMKIIα</sub> (N<sub>2</sub>) | t-test 2 | 1000 | 926 | neurons | <10<sup>-5</sup> |
|  | paAIP2<sub>SST</sub> (N<sub>2</sub>) | t-test 2 | 1000 | 397 | neurons | 0.0044 |
|  | paAIP2<sub>PV</sub> (N<sub>2</sub>) | t-test 2 | 1000 | 492 | neurons | <10<sup>-5</sup> |

**Table S2.**
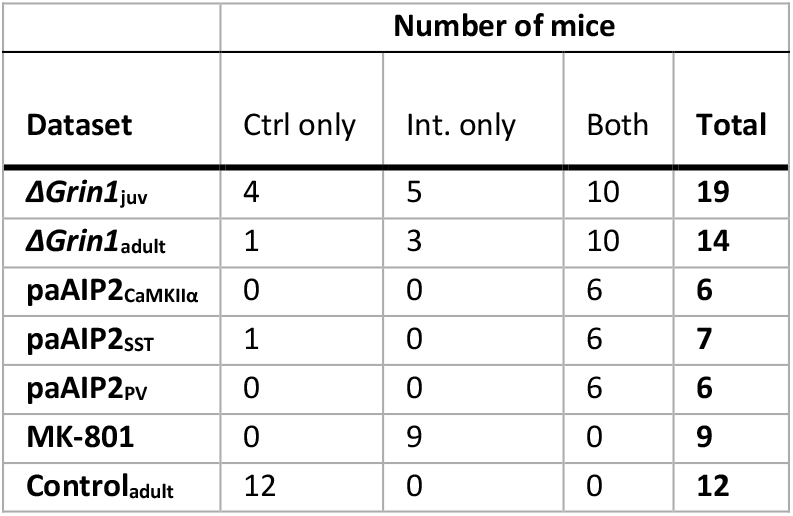
Number of mice per group. We imaged all experimental mice on both hemispheres whenever possible. The table below lists the number of mice as a function of whether they were imaged in the control (Ctrl) or the intervention (Int.) hemisphere only, or in both hemispheres.

## Data and code availability

Software for controlling the two-photon microscope and preprocessing of the calcium imaging data is available on https://sourceforge.net/projects/iris-scanning/. Raw data and code to generate all figures of this manuscript are available on https://data.fmi.ch/PublicationSupplementRepo/.

## Acknowledgements

We thank all the members of the Keller lab for discussion and support and Tingjia Lu and Daniela Gerosa for vector production. We thank Ryohei Yasuda for valuable advice and help with paAIP2. This project has received funding from the Swiss National Science Foundation, the Novartis Research Foundation, and the European Research Council (ERC) under the European Union’s Horizon 2020 research and innovation programme (grant agreement No 865617).

## Author contributions

FW designed and performed the experiments and analyzed the data. Both authors wrote the manuscript.

